# Long lifetime and selective accumulation of the A-type lamins accounts for the tissue specificity of Hutchinson-Gilford progeria syndrome

**DOI:** 10.1101/2023.02.04.527139

**Authors:** John Hasper, Kevin Welle, Kyle Swovick, Jennifer Hryhorenko, Sina Ghaemmaghami, Abigail Buchwalter

## Abstract

Mutations to the *LMNA* gene cause laminopathies including Hutchinson-Gilford progeria syndrome (HGPS) that severely affect the cardiovascular system. The origins of tissue specificity in these diseases are unclear, as the A-type Lamins are abundant and broadly expressed proteins. We show that A-type Lamin protein and transcript levels are uncorrelated across tissues. As protein-transcript discordance can be caused by variations in protein lifetime, we applied quantitative proteomics to profile protein turnover rates in healthy and progeroid tissues. We discover that tissue context and disease mutation each influence A-type Lamin protein lifetime. Lamin A/C has a weeks-long lifetime in the aorta, heart, and fat, where progeroid pathology is apparent, but a days-long lifetime in the liver and gastrointestinal tract, which are spared from disease. The A-type Lamins are insoluble and densely bundled in cardiovascular tissues, which may present an energetic barrier to degradation and promote long protein lifetime. Progerin is even more long-lived than Lamin A/C in the cardiovascular system and accumulates there over time. Progerin accumulation interferes broadly with protein homeostasis, as hundreds of abundant proteins turn over more slowly in progeroid tissues. These findings indicate that potential gene therapy interventions for HGPS will have significant latency and limited potency in disrupting the long-lived Progerin protein. Finally, we reveal that human disease alleles are significantly over-represented in the long-lived proteome, indicating that long protein lifetime may influence disease pathology and present a significant barrier to gene therapies for numerous human diseases.

**Significance statement:** Many human diseases are caused by mutations to broadly expressed proteins, yet disease mysteriously manifests only in specific tissues. An example of this is Hutchinson-Gilford progeria syndrome (HGPS), which is caused by a mutation to the Lamin A/C protein. We show that this mutation slows the turnover of Lamin A/C proteins in disease-afflicted tissues, causing the mutant “Progerin” protein to accumulate over time and interfere with the normal turnover of hundreds of other proteins. Because Progerin is a long-lived protein, effective therapies for this disease will need to attack the protein and not just the gene that encodes it.

## Introduction

The nuclear lamina is an intermediate filament meshwork that underlies the nuclear envelope. In mammals, this structure is composed of two classes of Lamin proteins: the “A-type” lamins (Lamins A and C), and the “B-type” lamins (Lamins B1 and B2). Lamins assemble into bundled filaments that strengthen the nucleus and scaffold the genome.

The *LMNA* locus encodes the Lamin A and C proteins and is a hotspot for disease-linked mutations; hundreds of mutations to *LMNA* have been linked to at least 15 distinct syndromes, collectively referred to as “laminopathies”(*1*). While the Lamin A/C proteins are broadly expressed, laminopathies primarily afflict the cardiovascular system, muscle, bone, and fat(*1*). Hutchinson-Gilford progeria syndrome (HGPS) is a rare and devastating autosomal dominant laminopathy that resembles physiological aging at an accelerated rate, with rapid deterioration of the skin, adipose tissue, and cardiovascular system. Most cases of HGPS are caused by a single base pair substitution (GGC > GGT) that activates a cryptic splice site, resulting in the in-frame deletion of a portion of exon 11 in *LMNA* and the production of a toxic, dominant negative “Progerin” protein.

Several gene therapy approaches have recently been tested in mouse models of HGPS, ranging from disrupting the mutant allele with CRISPR/Cas9 indels(*2, 3*), correcting the mutant allele with base editing(*4*), or interfering with the unique RNA splicing event that produces the *Progerin* transcript(*5–8*). Because HGPS is caused by the toxic effects of a dominant negative protein, an effective gene therapy will need to induce the removal and replacement of as much of the Progerin protein as possible. However, recent work indicates that gene therapies have variable effects on Progerin protein levels *in vivo*; for instance, in various treatment regimes that effectively clear the mutant protein from the liver, the protein persists to some extent in the cardiovascular system (*3–6, 8, 9*). In one particularly striking case, high levels of Progerin were found to persist in the cardiovascular system after over 5 months of treatment with splice-interfering antisense oligonucleotides, long after effective depletion of the *Progerin* transcript had been achieved (*6*). Altogether, these findings indicate that removing Progerin protein is a major bottleneck that may limit the potency of HGPS gene therapies.

Why is Progerin so intractable to removal in the cardiovascular system? Progerin could be produced in particularly high amounts in cardiovascular tissues, which are known to express high levels of Lamin A/C(*10*). A second, non-exclusive possibility is that Progerin is especially long-lived in the cardiovascular system. Notably, the closely related B-type Lamin proteins have extremely long lifetimes of months to years in the murine nervous system(*11, 12*). The lifetimes of the A-type Lamins within tissues have not been defined. We previously profiled protein lifetimes in cultured progeroid fibroblasts and found no evidence that the HGPS-causative mutation impairs the turnover of Lamin A/C(*13*). However, these experiments could not address the potential influence of tissue context on the turnover rate of Lamin A/C and/or Progerin.

Here, we deploy our recently developed dynamic proteomic approach, turnover and replication <u>a</u>nalysis by isotope labeling (TRAIL)(*14*) to define the turnover rate of the Lamin A/C and Progerin proteins within tissues of healthy and progeroid mice. We report that Lamin A/C turns over an order of magnitude more slowly within disease-afflicted tissues compared to disease-spared tissues. Further, we show that the Progerin mutant is even more long-lived than wild-type Lamin A/C in the cardiovascular system, leading us to estimate a months-long lifetime of this toxic mutant protein in these diseased tissues. In contrast, we show that protein abundance does not correlate with disease susceptibility in human or mouse tissues. By evaluating the incidence of human disease alleles across protein turnover deciles, we reveal that human disease alleles are generally over-represented in the long-lived proteome. These findings have major implications for both the pathogenesis and treatment of laminopathies and establish a novel paradigm for rare human diseases caused by the action of long-lived mutant proteins.

## Results

### High abundance of Lamin A/C/Progerin proteins is not restricted to cardiovascular tissues

High Progerin abundance could both drive disease and thwart gene therapies. We reasoned that tissues that express high levels of Lamin A/C in healthy individuals may also express more Progerin in diseased individuals, and we mined a quantitative proteomic atlas of 29 human tissues(*15*) to systematically evaluate the relationship between Lamin A/C protein expression and laminopathy pathology. This tissue proteome atlas relies on intensity-based absolute quantification (iBAQ) to accurately quantify protein abundance(*16*) and on the histone-based “proteomic ruler” approach(*17*) to estimate protein copy numbers per cell. These data indicate that the human Lamin A/C proteins are extremely abundant in smooth muscle (Fig. 1A), consistent with previous reports in mouse tissues(*18, 19*). Because vascular smooth muscle is prominently affected in *LMNA*-linked Hutchinson-Gilford progeria syndrome(*20, 21*), these data could suggest that high Lamin A/C abundance correlates with disease vulnerability. However, the heart and adipose tissue are also afflicted in HGPS and other laminopathies, but these tissues express Lamin A/C at a level comparable to many unaffected human tissues (Fig. 1A). In fact, 26/29 of the analyzed tissues are estimated to produce more than 10^7^ copies of Lamin A/C per cell (Fig. S1A), indicating that high expression levels of Lamin A/C are not sufficient to drive disease.

**Figure 1.**
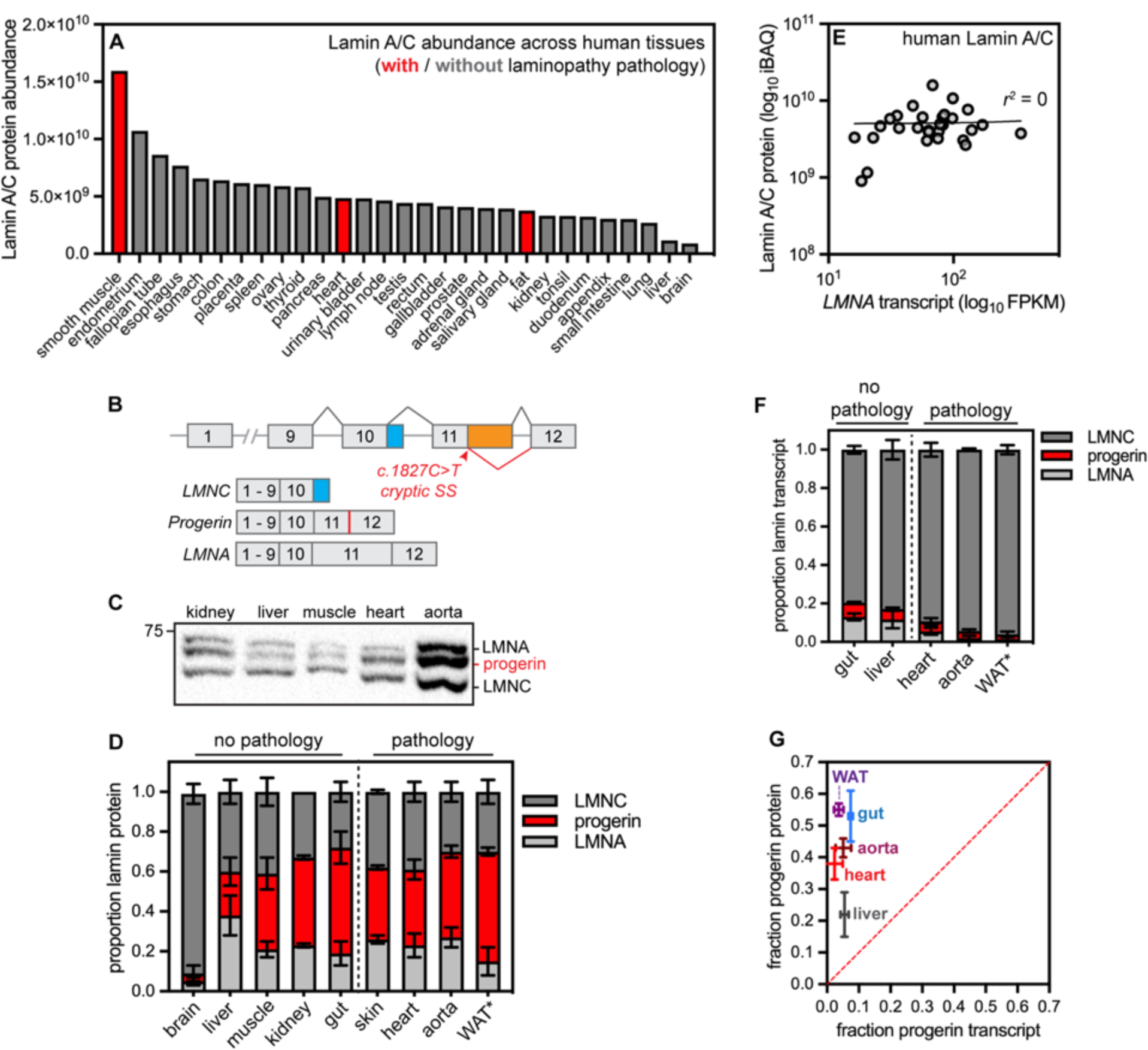
Poor correlation between trasncript and protein abundance for the long-lived disease-linked protein Lamin A/C. (A) iBAQ quantification of Lamin A/C absolute protein abundance across 29 human tissues. Data reanalyzed from Wang et al., *Mol Syst Biol* 2019. (B) Diagram of the *LMNA* locus, which produces the *LMNA* (exons 1-12) and *LMNC* (exons 1­10) transcripts. In HGPS, a base pair substitution activates a cryptic splice site, leading to the production of the toxic *Progerin* transcript that contains an in-frame deletion in exon 11. (C) Representative Western blot of tissue extracts from 9-month old *LMNA^G6O9G/t^* mice. 20 ug of protein loaded per lane. (D) Quantification of proportional Lamin A, Lamin C, and Progerin protein isoform abundance by Western blotting and densitometry in 9-month old *LMNA^GmG/^\** mice, ‘indicates that WAT was analyzed from 3-month-old animals because of the rapidly progressing lipodystrophy in these mice. (E) Lamin A/C absolute protein abundance (iBAQ) and *LMNA* RNA abundance (RNAseq FPKM) are very poorly correlated (r^2^ = 0) across 29 human tissues. Data reanalyzed from Wang et al„ *Mol Syst Biol* 2019. (F) Proportional abundance of *LMNA, LMNC,* and *Progerin* transcript abundance determined by digital droplet PCR in tissues from 9-month old *LMNA^G6mG/^’* mice, ‘indicates that WAT was analyzed from 3-month-old animals. (G) Proportional abundance of *Progerin* transcript and Progerin protein are very poorly correlated across tissues from *LMNA^G609G^’’* mice.

The abundance ratio of the A-type (Lamin A/C) to B-type (Lamin B1 and B2) lamins influences nuclear mechanics(*18*) and it has been proposed that tissues with a high ratio of A-type:B-type lamins might be most sensitive to *LMNA* mutations(*19*). Consistent with previous analyses in the mouse(*19*), the ratio of A-type:B-type lamins is very high in the human heart (8.6:1), smooth muscle (7.2:1), and fat (4.3:1) which are especially vulnerable to *LMNA* mutations (Fig. S1D). However, this ratio is also comparably high in many unaffected tissues (Fig. S1D). Therefore, a high ratio of A-type to B-type Lamins is not sufficient to drive HGPS.

We next considered the alternative possibility that Progerin abundance varies independently of Lamin A/C abundance across diseased tissues. The genetic lesion that causes HGPS activates a cryptic splice site, resulting in the in-frame deletion of a portion of exon 11 in *LMNA* without altering the shorter *LMNC* isoform (Fig. 1B). This splice site operates with incomplete efficiency, such that both *Progerin* and wild type *LMNA* transcripts can be produced from the mutant allele; variable splice site use could thus modulate *Progerin* production across tissues(*22*). To explore this possibility, we turned to a genetically modified mouse model that harbors the HGPS-causative C>T substitution in the endogenous mouse *LMNA* locus (*LMNA^G609G/+^)*(*5*).

We evaluated the proportional abundance of the Lamin A, Lamin C, and Progerin proteins by Western blotting across tissues in wild type (Fig. S2) and progeroid mice (Fig. 1C-D) by detection with an antibody that detects an epitope shared across all A-type lamins (see Methods). We reasoned that this approach was the most relevant way to quantify the dose of the toxic Progerin protein, as the ratio of Progerin to other wild type Lamin isoforms has been proposed to influence pathology(*19*). We found that Progerin contributes significantly to the total amount of A-type Lamins detected in many tissues - up to 50% in some cases(Fig. 1C-D). Puzzlingly, however, the proportional abundance of Progerin does not correlate with HGPS pathology; the disease variant is highly abundant in afflicted tissues, such as the aorta and adipose tissue, as well as in tissues that remain functional, such as the kidney and large intestine. Altogether, these analyses indicate that protein abundance alone cannot explain either the vulnerability of specific tissues to *LMNA* mutations or the resistance of Progerin to therapeutic interventions.

### Lamin A/C/Progerin protein and transcript levels are very poorly correlated across tissues

What other factors could modulate both the responsiveness of Progerin to treatment and the effects of Progerin on a tissue? One potential influence could be the lifetime of the Lamin A/C/Progerin proteins. A protein’s lifetime reflects its rates of synthesis and degradation; short-lived proteins are frequently renewed, while long-lived proteins can persist for weeks or even months within mammalian tissues(*11*). Because long-lived proteins are rarely synthesized and rarely degraded, their abundance may be poorly sensitive to changes in their transcript levels. To explore the relationship between *LMNA* transcript and Lamin A/C protein abundance, we compared tissue-matched RNAseq and proteomic data across 29 human tissues(*15*). This analysis revealed that *LMNA* transcript and Lamin A/C protein abundances are uncorrelated (Fig. 1E, *r^2^* ∼ 0). For comparison, the enzyme SYK has a high correlation between transcript and protein levels across tissues(*15*) (Fig. S3, *r^2^*= 0.89). We next used digital droplet PCR to precisely quantify the abundance of the *LMNA, LMNC,* and *Progerin* transcripts in progeroid mouse tissues and observed that *Progerin* makes up a minor proportion of the transcripts produced from the *LMNA* locus (Fig. 1F), yet these same tissues produce high levels of Progerin protein (Fig. 1C-D). The proportional abundance of *Progerin* transcript and Progerin protein are thus very poorly correlated (Fig. 1G). Taken together, these data indicate that Lamin A, Lamin C, and Progerin protein levels are post-transcriptionally modulated in human and mouse tissues. Similarly poor correlations between transcript and protein levels have been observed for extremely long-lived proteins, such as components of the nuclear pore complex(*23*), leading us to speculate that Lamin A/C and/or Progerin are also long-lived proteins.

### Lamin A/C/Progerin are long-lived proteins in the cardiovascular system

To test the hypothesis that the A-type Lamins have long lifetimes, we sought to determine the turnover rates of these proteins within the tissues of healthy and progeroid mice. Proteome-wide quantification of protein stability can be achieved *in vivo* by supplying mice with chow containing the stable, non-toxic isotope ^15^N, then tracking the rate of incorporation of ^15^N-labeled amino acids into the proteome over time by mass spectrometry(*14, 24*). We recently used ^15^N metabolic labeling to profile trends in protein turnover across healthy mouse tissues(*14*), and here applied the same 6-timepoint, 32-day labeling timecourse approach to age-matched *LMNA^G609G/+^* mice. These mice exhibit weight loss, bone abnormalities, atherosclerosis, and cardiac dysfunction before succumbing to the disease between 9 and 12 months of age(*5*). We focused our analyses on young adult mice at early stages of disease (9 to 12 weeks of age), and quantified protein lifetimes in tissues that exhibit progeroid pathology (the aorta, heart, and white adipose tissue) as well as tissues that are spared from disease (the intestine and liver). We defined protein turnover rates (*k_t_*) and corresponding predicted half-life (*t_1/2_*) values for 2744 proteins in the large intestine, 2098 proteins in the liver, 1881 proteins in the aorta, 1429 proteins in the white adipose tissue, and 1608 proteins in the heart (Fig. S4A-B; Table S1). Consistent with our previous analyses in healthy tissues, proteins generally turn over more rapidly in the progeroid intestine (median *t_1/2_* 1.6 days) and liver (median *t_1/2_* 2.7 days) than in the progeroid aorta (median *t_1/2_* 3.8 days), heart (median *t_1/2_* 7.1 days) or fat (median *t_1/2_* 9.2 days).

While the goal of these experiments is to determine whether the A-type Lamins exhibit variable lifetimes across tissues, a major confounding factor limits our ability to interpret differences in protein lifetimes across tissues *in vivo*: the influence of cell turnover. That is, protein turnover *in vivo* is shaped both by selective proteolytic degradation and non-selective dilution during cell turnover. We recently developed a method, turnover and replication <u>a</u>nalysis by ^15^N isotope labeling (TRAIL) (Hasper et al 2022), to quantify both protein and cell lifetimes from the same ^15^N-labeled tissues to determine cell turnover-corrected protein degradation rates (Fig. 2A). We applied this method to determine cell turnover rates (*k_div_*) and doubling times within progeroid intestine, liver, fat, and heart (Fig. S4C-D; Table S2). (Cell turnover was not determined for the aorta due to limited sample.) Overall, cell doubling times were similar in progeroid and healthy tissues with one exception: the white adipose tissue, where progeroid cells had a doubling time of ∼55 days while age-matched wild type cells had a doubling time of ∼78 days. This observation likely reflects increased proliferation and/or increased death of adipocytes, which are the most abundant cell type found in fat depots. Notably, over-expression of Progerin blocks the terminal differentiation of adipocyte precursors(*25*), and HGPS patients suffer from severe lipodystrophy(*26*). We speculate that *LMNA^G609G/+^* mice are in the early stages of lipodystrophy at 9 weeks of age, as these animals are indistinguishable in weight compared to their littermates(*5*) (Fig. S5), although their fat depots are moderately decreased in size.

**Figure 2.**
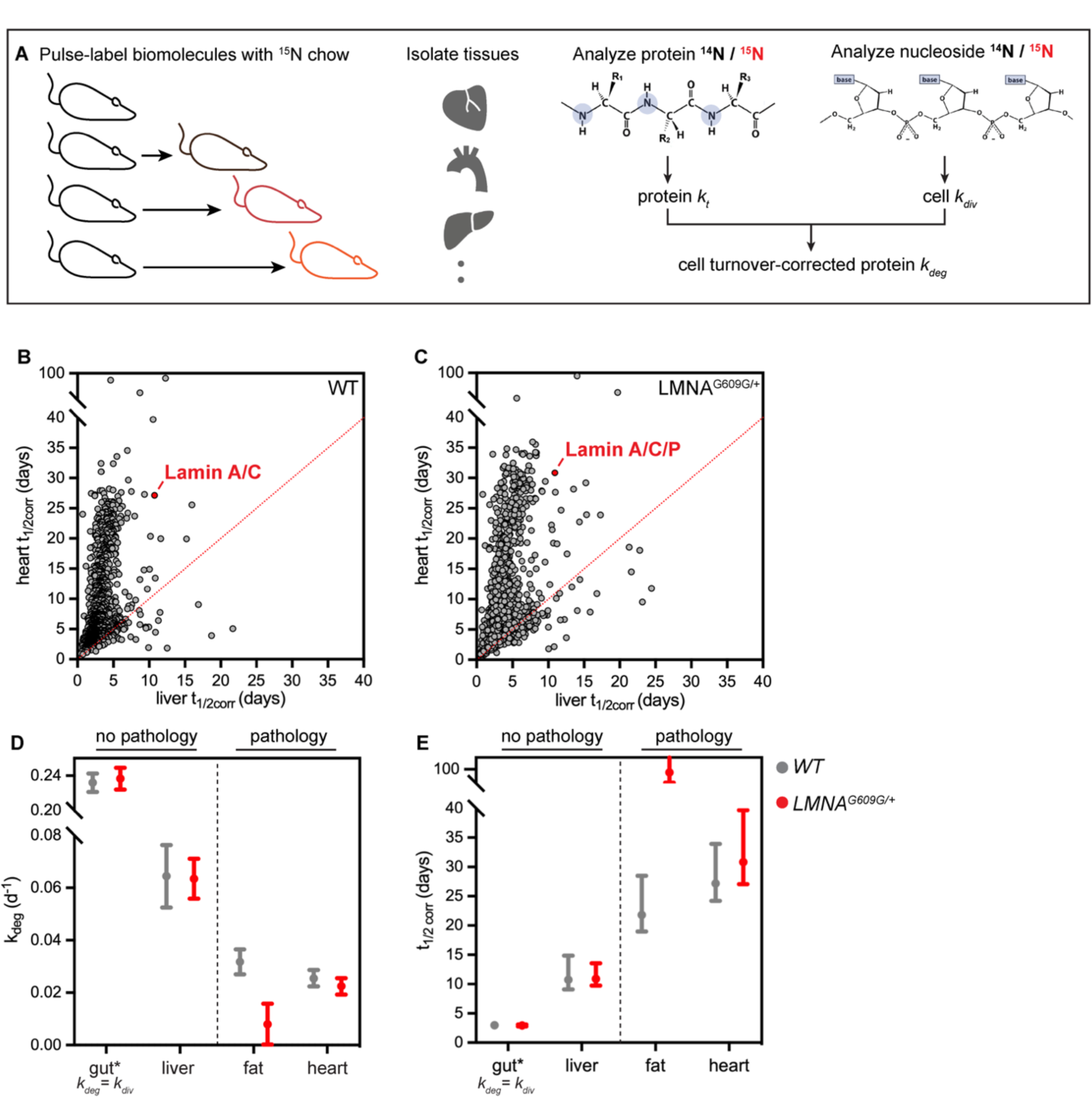
(A) Diagram of turnover and replication by ^15^N isotope labeling (TRAIL) method for determining protein turnover (k_t_) and cell turnover (k_dlv_) by tracking ^16^N incorporation into newly synthesized protein and genomic DNA, respectively. These two parameters are used to determine average bulk cell cycle-corrected protein degradation rates (k_deg_). (B-C) Distribution of half-lives for proteins detected in both the liver and heart of healthy (B) and *LMNA^GS09G/^\** (C) mice. The A-type Lamin proteins (red) are among a group of outliers with a dramatically longer lifetime in the heart than in the liver. (D-E) Cell cycle-corrected protein degradation rates (k_deg_) (D) and half-lives (t„_2corr_) (E) for A-type Lamin proteins in tissues of healthy (gray) and *LMNA^a60SG/^\** (red) mice. * denotes that A-type Lamin turnover is equivalent to cell turnover rate in the gut.

By subtracting cell turnover rates (*k_div_*) from protein turnover rates (*k_t_*), we extrapolated cell cycle-corrected protein degradation rates (*k_deg_*) and predicted half-lives *(t_1/2corr_*) for proteins in the progeroid intestine, liver, heart, and fat. As we previously observed in wild type animals(*14*), protein turnover rates vary widely across tissues even after cell cycle correction (Fig. 2B-C; Fig. S4F; Table S1). We infer that the rate of cell division as well as environmental factors such as local metabolism or the activity and selectivity of proteostasis networks shape protein lifetimes across tissues(*14*).

We were able to determine the average turnover behavior of the Lamin A, Lamin C, and Progerin proteins. We find that the A-type lamins have highly variable lifetimes across both healthy and progeroid tissues. The distribution of half-lives for proteins detected in both the liver and heart tissue of wild type (Fig. 2B) and progeroid (Fig. 2C) animals illustrates this point, as the A-type lamins are among a small group of outlier proteins that have a dramatically longer lifetime in the heart than in the liver. Overall, the lifetimes of the A-type lamins are longer in tissues that exhibit progeroid pathology (the aorta, heart, and fat) compared to those that are spared from pathology (the liver and intestine) in both wild type and progeroid animals (Table S1). After using TRAIL to quantify cell turnover rates and determine corrected protein turnover rates, the differences between A-type lamin turnover flux across these tissues were even more stark (Fig. 2B-C; cell turnover not determined for the aorta due to limited sample); indicating that these proteins turn over dramatically more slowly in the fat and the heart. After cell cycle correction, the predicted half-life for Lamin A/C in the healthy heart and fat is approximately 3 weeks. Further, the average turnover rate of the closely related Lamin A, Lamin C, and Progerin proteins is slowed to >1 month in the progeroid heart and fat compared to healthy tissues, raising the possibility that the mutant Progerin protein turns over even more slowly than wild type Lamin A/C (Fig. 2E).

### Biochemical influences on Lamin A/C and Progerin protein lifetime

Why are the A-type Lamins readily turned over in the liver, but more slowly turned over in the heart? The lamins form head-to-tail filaments that bundle laterally(*27*) and stiffen(*28*) or undergo structural changes(*29*) in response to mechanical force. Because the heart is a contractile tissue under high strain, we hypothesized that differences in the folding and/or assembly state of the Lamin proteins in the heart might underlie the proteins’ longer lifetime in this tissue. To test this hypothesis, we subjected tissue homogenates to sequential extractions of increasing stringency - using salt, detergent, and denaturants(*30*) – and compared the extent of Lamin A/C/Progerin extraction after each step. These analyses indicated that the A-type lamins are poorly extractable in heart tissue, where complete extraction was possible only in strongly denaturing conditions (4M urea) (Fig. 3A-B). In the liver, in contrast, the A-type lamins were efficiently extracted by treatment with high salt (Fig. 3C-D). These observations strongly imply physicochemical differences in the assembly state of the Lamin polymers that correlate with the differences in protein turnover rate between these tissues.

**Figure 3.**
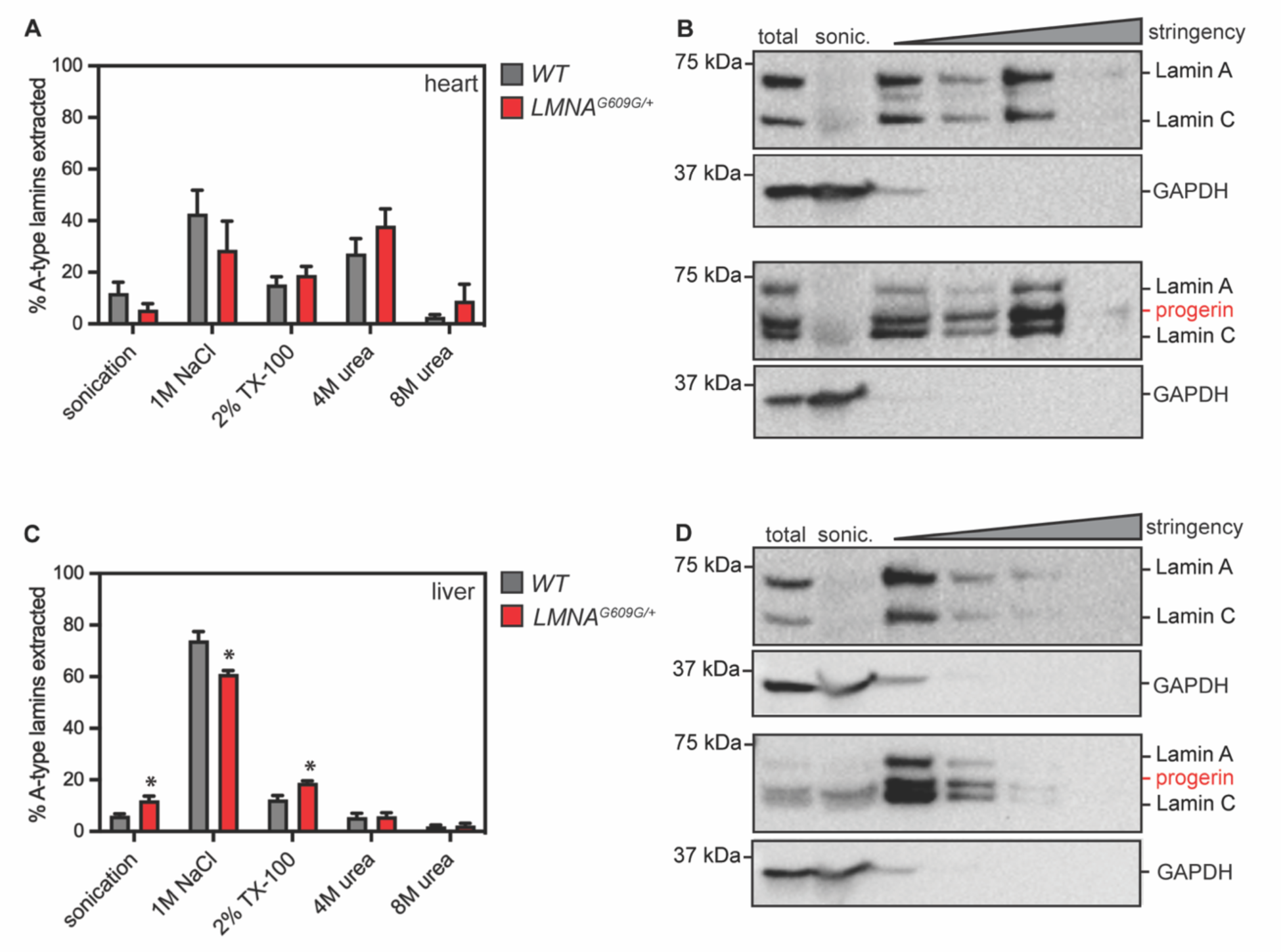
Serial extraction of LAmin A/C/Progerin proteins from heart (A-B) or liver (C-D) tissue of 4-month-old healthy and *LMNA^G609G/^\** mice. GAPDH indicates effective lysis and reelase of cytosol in each tissue. While Lamin A/C/Progerin are readily extractable with high salt in the liver (C-D) strong dénaturants are required to completely extract these proteins from the heart (A). * indicates significant differences in protein solubility between wild type and *LMNA^G609G^’\** liver tissue.

### Progerin exhibits impaired turnover and accumulates in the cardiovascular system

Progerin retains a hydrophobic lipid modification that is only transiently present on wild-type Lamin A(*31*), raising the possibility that this disease variant could be more aggregation-prone and/or more recalcitrant to turnover than the normal protein. Lamin A is synthesized as a pre-protein that matures through a series of post-translational modifications. These include a lipid modification (farnesylation) followed by proteolytic cleavages that remove the lipid-modified C-terminus to generate mature Lamin A(*32*). The HGPS mutation deletes a key protease cleavage site(*31*), generating the constitutively farnesylated Progerin protein. We could not determine whether these closely related isoforms exhibit distinct turnover rates by shotgun proteomics (Fig. 2), as the small subset of peptides that distinguish Lamin A, Lamin C, or Progerin from each other were not detected.

To determine whether the HGPS mutation extends protein lifetime, we immunoprecipitated the A-type lamins from metabolically labeled progeroid heart, aorta, and liver and performed parallel reaction monitoring mass spectrometry (PRM-MS)(*33*) to quantify peptides unique to Lamin A *versus* Progerin (Fig. 4A-B; Table S3). We did not monitor Lamin-C-only peptides, as Lamin C is distinguishable from Lamin A only at its C-terminal 6 amino acids, and this region does not generate any quantifiable and unique tryptic peptides. For maximum sensitivity, we focused our analyses on the labeling timepoint that was closest to the predicted half-life of the A-type lamins in each tissue: 32 days for the heart, 16 days for the aorta, and 8 days for the liver. By quantifying the ^15^N/^14^N isotope abundance ratio for individual peptides unique to each protein isoform, we could compare the relative extent of turnover. These analyses revealed that Progerin turns over significantly more slowly than wild type Lamin A in the heart (Fig. 4C-E) and aorta (Fig. 4F-H) but not in the liver (Fig. 4I-K). Comparing these isoform-resolved data to our bulk A-type lamin turnover data (Table S3) leads us to estimate that Progerin’s lifetime within the cardiovascular system is on the order of months. Altogether, these protein turnover analyses indicate that both tissue context and disease mutations modulate the lifetime of the A-type lamin proteins.

**Figure 4.**
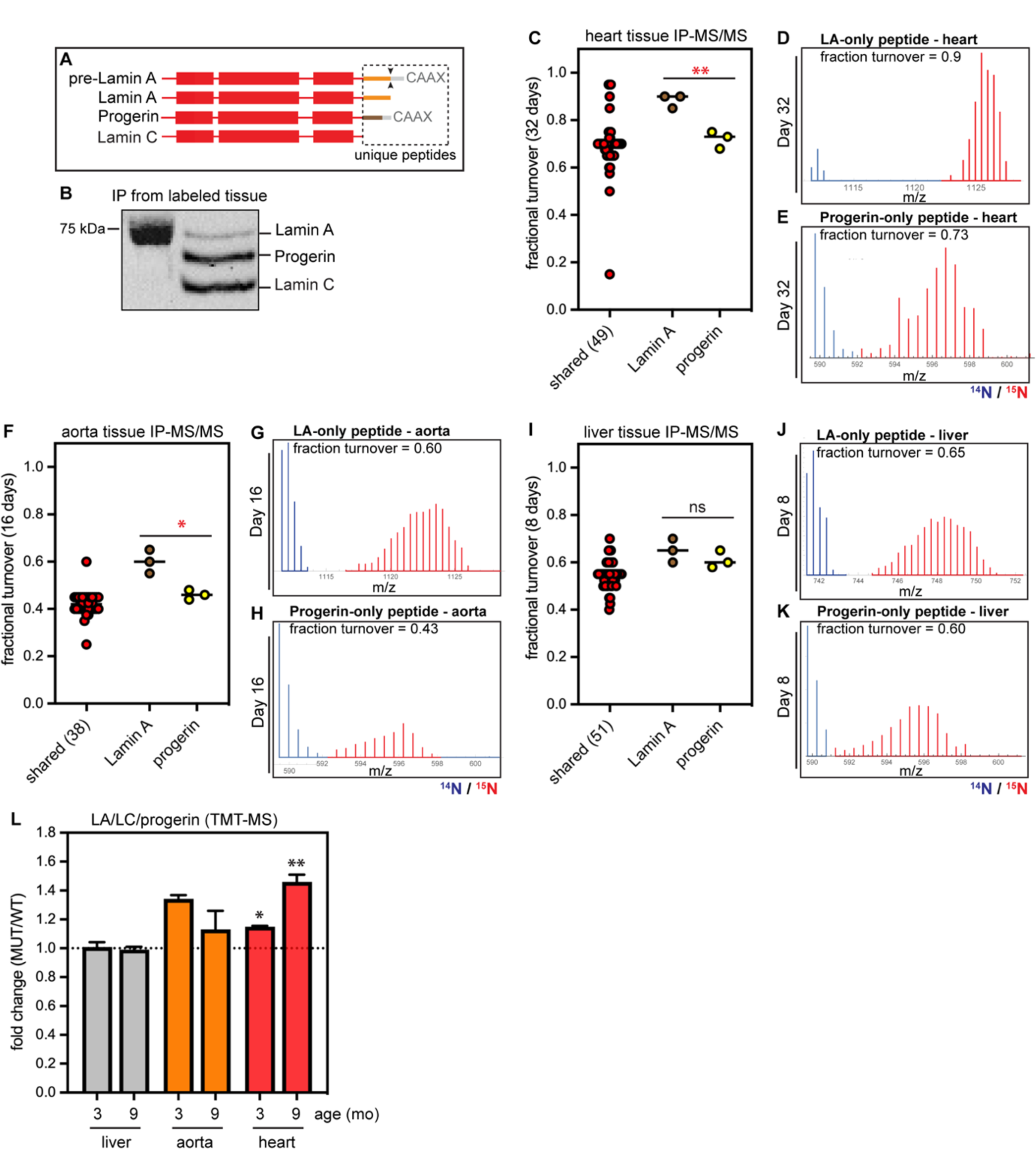
(A) Diagram of shared versus unique regions of the pre-Lamin A, Lamin A, Progerin, and Lamin C protein isoforms. Dashed box, region containing unique peptides. (B) Representative immunoprecipitation of Lamin A/C/Progerin from ^15^N-labeled mouse tissue. (C) PRM-MS detect-ion of a Lamin A-specific peptide (brown), Progerin-specific peptide (yellow), and shared peptides (red) In heart tissue after 32 days of 15N labeling. ** indicates that label incorporation is significantly slower in Progerin than Lamin A, Indicating slower turnover. (D-E) Representative spectra of Lamin A-specific peptide (D) and Progerin-specific peptide in the heart, “old” "’N-labeled peptide shown In blue and “new" ^15^N-labeled peptide shown in red. (F-K) PRM-detection of Lamin A and Progerin-specific peptides in the aorta (F-H) and liver (l-K). Progerin turns over more slowly in the aorta (*) but not in the liver (ns). (L) Analysis of relative protein abundance In wild-type versus *LMNA^SS09S^’\** mouse tissues at 3 months and 9 months of age by TMT-MS. Total Lamin A/C/Progerin abundance increases in the progerold aorta and heart, but not in the progeroid liver.

If Progerin turnover is impaired compared to wild type Lamin A, one might predict that Progerin would accumulate over time. We used tandem mass tagging (TMT) and proteomics(*34*) to quantify relative protein abundance in healthy *versus* progeroid tissues from young adult mice (3 months old) and from animals nearing the end of the HGPS mouse lifespan (9 months old). These data indicated that the total dose of A-type lamin proteins is significantly increased in the progeroid heart, but not the liver; while the dose of these proteins trends upward in the diseased aorta, this trend is not significant due to variability across samples (Fig. 4L). Because this analysis cannot distinguish Progerin from Lamin A, these data could indicate either that Progerin accumulates over time, or that Progerin also interferes with the turnover of wild-type Lamin A, leading to accumulation of all A-type lamin isoforms in diseased tissue.

### Progerin expression impairs global proteostasis

Our data indicate that the Progerin disease variant accumulates selectively within the cardiovascular system over time. We hypothesized that as Progerin accumulates within diseased tissue, it could sequester other proteins and interfere with their turnover. Consistent with this hypothesis, we observed global changes in protein stability in progeroid tissues (Fig. 5A). At an individual protein level, 24-40% of quantified proteins have significantly extended lifetimes in progeroid tissues compared to healthy tissues (Fig. 5B). The magnitude of this effect is most pronounced in the heart and adipose tissue, where this subset of proteins experience a median lifetime extension of approximately 2 days compared to age-matched wild-type animals (Fig. 5C). These proteins are constituents of various protein structures including mitochondria, ribosomes, the nucleus, and the cytoskeleton (Fig. 5E-G). As these proteins are components of such diverse cellular structures, we hypothesized that shared physicochemical features, not protein localization, might underlie their sensitivity to Progerin expression. Indeed, we found that Progerin-sensitive proteins are much more abundant than Progerin-insensitive proteins (Fig. 5D). Altogether, the ability of Progerin to impair the turnover of abundant proteins associated with a broad range of cellular structures is consistent with an aggregation-based mechanism of toxicity, as highly abundant proteins are more vulnerable to aggregation(*35, 36*)

**Figure 5.**
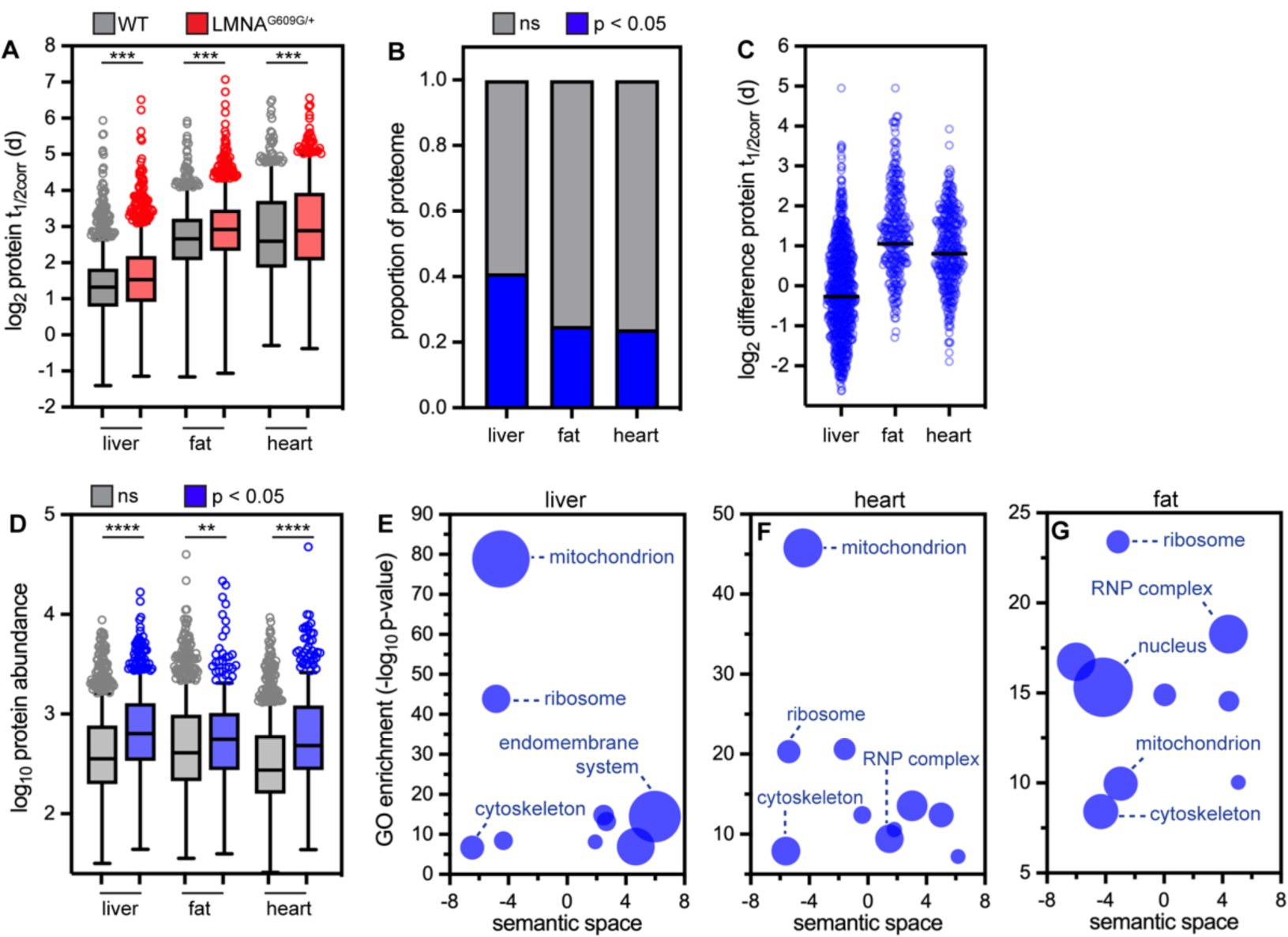
(A) Analysis of distribution of human disease alleles (OMIM) across deciles of protein turnover rates in healthy mouse tissues. Map color indicates significance of over-representation (yellow) or under-representation (blue) of disease alleles in each decile. ** indicates that human disease alleles are significantly over-represented in the 1st, 2nd, and 3rd deciles of k_deg_rate (most long-lived) in the heart; in the 1st decile in the fat; and in the 2nd decile in the liver (Fisher’s exact test). Table II shows protein turnover rate data for selected long-lived proteins linked to human disease. Data re-analyzed from Hasper et al., BioRXiV 2022. See Table S4

**Figure 5.**
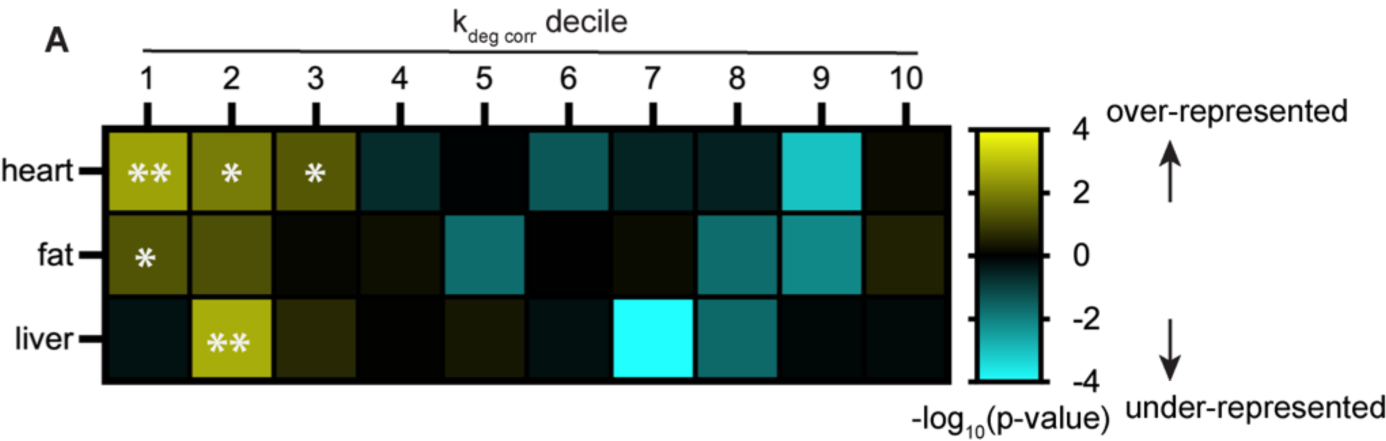
(A) Predicted protein half-lives determined for proteins in liver, fat, and heart muscle of 3-month-old healthy and LMNA^G^“^9G/^* mice. *** indicates a significant decrease in protein turnover flux in progeroid tissues. Data from wild type animals reproduced from Hasper et al., *Mol Syst Biol* 2023. (B) Proportion of proteins in each tissue with significantly different turnover rates in progeroid animals versus wild type animals (t-test). (C) Difference in protein lifetime (MUT-WT) for proteins with significantly different lifetimes in progeroid animals versus wild type animals. Liver median difference = + 0.8 days (n = 841); fat median difference +2.1 days (n = 349); heart median difference +1.75 days (n = 373). (D) Comparison of protein abundance between proteins with significantly altered lifetime in progeroid tissue (blue) versus proteins with unchanged lifetime in progeroid tissue (gray). *“* indicates that proteins with impaired turnover in progeroid tissue are more abundant than proteins with un­changed turnover in progeroid tissue (Mann-Whitney test). (E-G) Subsets of Gene Ontology (GO) terms (cellular component) over-represented in proteins with significantly slower turnover in progeroid tissues compared to healthy tissues. Redundant GO terms were removed and non-redundant GO terms organized based on semantic similarity with REViGO. Bubble size corre­sponds to number of proteins associated with the GO term, ranging in size from 12 to 175 (E), 14 to 95 (F), and 10 to 157 (G).

### Human disease alleles are over-represented in the long-lived proteome

Our data suggest that protein lifetime modulates the effects of disease-causative mutations to cause tissue-specific disease. To explore the relationship between protein lifetime and human disease more systematically, we evaluated the distributions of proteins linked to human disease across our recently acquired datasets of protein turnover rates in healthy mouse heart, liver, and adipose tissues(*37*). We quantified the turnover rates of ∼1600 to ∼2100 proteins per tissue, and found that mutations in ∼600 to ∼800 of those proteins are linked to human disease in the OMIM database. Evaluating the distributions of these disease-linked proteins across turnover rate deciles revealed that proteins linked to human disease are significantly over-represented in the most long-lived proteins in each tissue – with the most dramatic over-representation apparent in the heart (Fig. 5A, Table I, Table S4). Selected disease-linked proteins found in the 1^st^ decile of turnover rates in heart, liver, or fat are shown in Table I (see Table S4 for complete list). In addition to *LMNA,* mutations to *CAV3, ACTA1, ACTC1, DYSF,* and *DSG2* are linked to cardiomyopathies and encode proteins that are long-lived in the heart. Some of these proteins, such as ACTA1 and ACTC1, are selectively expressed in cardiac muscle, which provides an obvious explanation for the tissue-specific consequences of mutations. However, similarly to Lamin A/C, the DYSF, DSG2, and CAV3 proteins are expressed in several tissues but are only long-lived in the heart (Fig. S6C-E), while the proteins KRT18 and AGL are broadly expressed but are only long-lived in the liver (Fig. S6F-G), where their mutations cause disease. These analyses overall reveal an unexpected enrichment of human disease alleles in extremely long-lived proteins and suggest that protein lifetime may influence disease etiology not just in laminopathies but also in other monogenetic human diseases.

**Table I.**
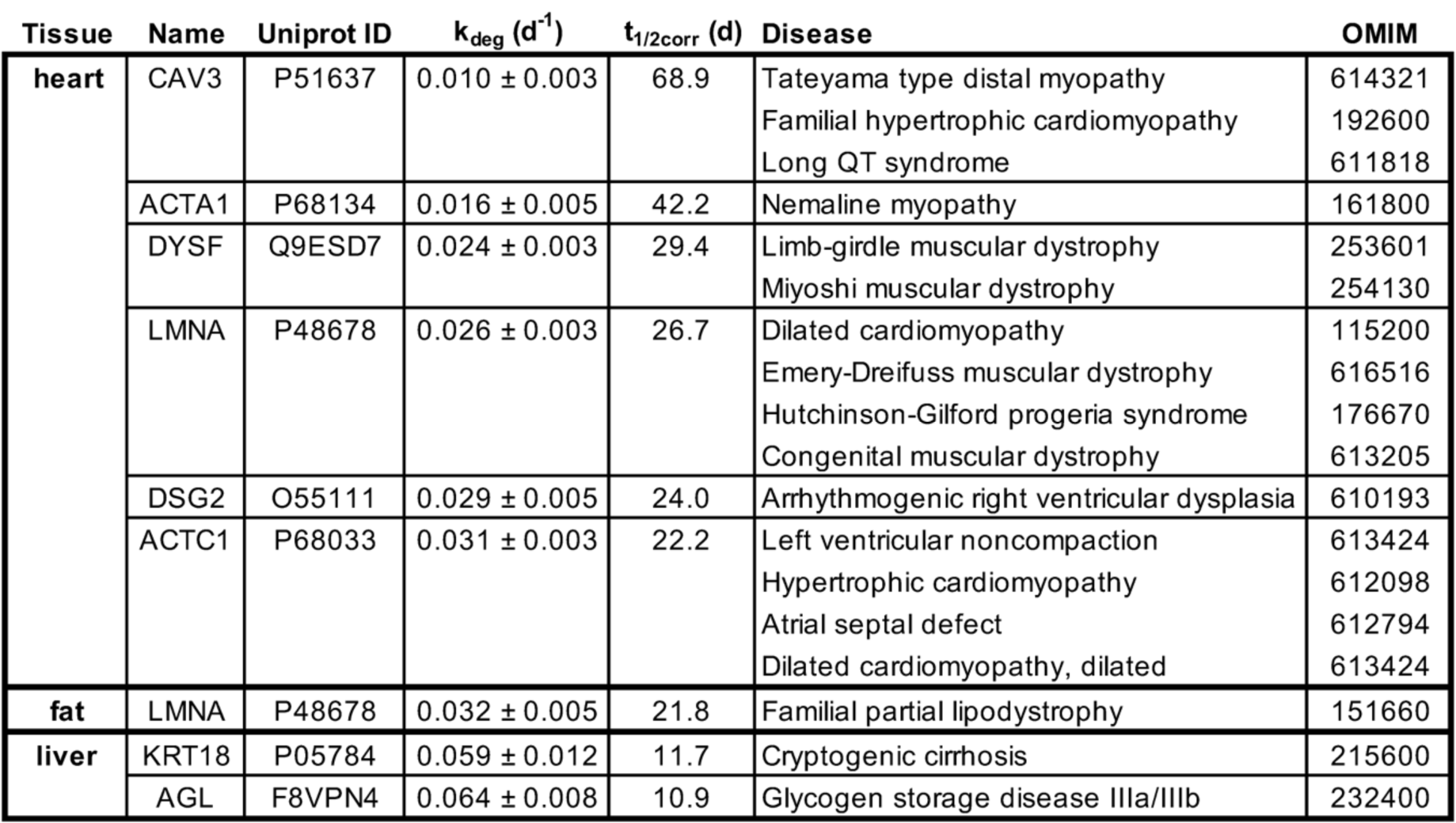
Long-lived proteins and human disease.

## Discussion

In this study, we applied quantitative proteomic approaches to explore the origins of tissue specificity in laminopathy syndromes, which are caused by mutations to the abundant and broadly expressed Lamin A/C protein. We determined that the Lamin A/C proteins have significantly longer lifetimes within the cardiovascular system and fat, which are disrupted in laminopathies, while these proteins have shorter lifetimes in disease-spared tissues such as the liver and intestine. We demonstrated that the A-type Lamins are more densely bundled and less extractable in the heart than in the liver. Since substrate unfolding is the major rate-limiting step in protein degradation(*38*), more densely bundled Lamin polymers may be more recalcitrant to protein turnover than the same proteins in a less densely bundled state. In the future, it will be important to explore how Lamin assembly state modulates the functions of the nuclear lamina.

Our findings suggest that HGPS and other laminopathies may arise as a consequence of disrupted proteostasis. In some tissues (such as the liver), the A-type Lamins may be maintained in a folded and functional state by consistent degradation and replacement (Fig. 7). In tissues where the Lamins are long-lived (such as the heart), in contrast, protein synthesis and degradation happen rarely. In this context, these long-lived proteins may undergo molecular “aging” – meaning that they may accumulate damage, misfold, and perhaps irreversibly aggregate in a time-dependent manner. Laminopathy-linked mutations may increase the propensity for misfolding and/or irreversible aggregation over time, leading to the tissue-specific and time-dependent accumulation of dysfunction. We found that the turnover of the HGPS-causative Progerin mutant is selectively impaired within the cardiovascular system, leading to its accumulation over time (Fig. 4). By analogy to neurodegenerative diseases caused by protein aggregation(*39*), Progerin aggregates could exert broad and pleiotropic influence on protein quality control by inducing co-aggregation of other proteins and/or by acting as a sink for the proteostasis machinery. Indeed, we observe impaired turnover of hundreds of abundant proteins in progeroid tissues (Fig. 5). These proteins include constituents of mitochondria, ribosomes, the nucleus and the cytoskeleton, and dysfunction related to each of these cellular structures has been reported in HGPS(*13, 40–42*). Decreased proteasome activity has been reported in fibroblasts derived from HGPS patients(*13, 43*), leading us to speculate that decreased flux through the proteasome could contribute to extension of protein lifetimes that we observe.

**Figure 7.**
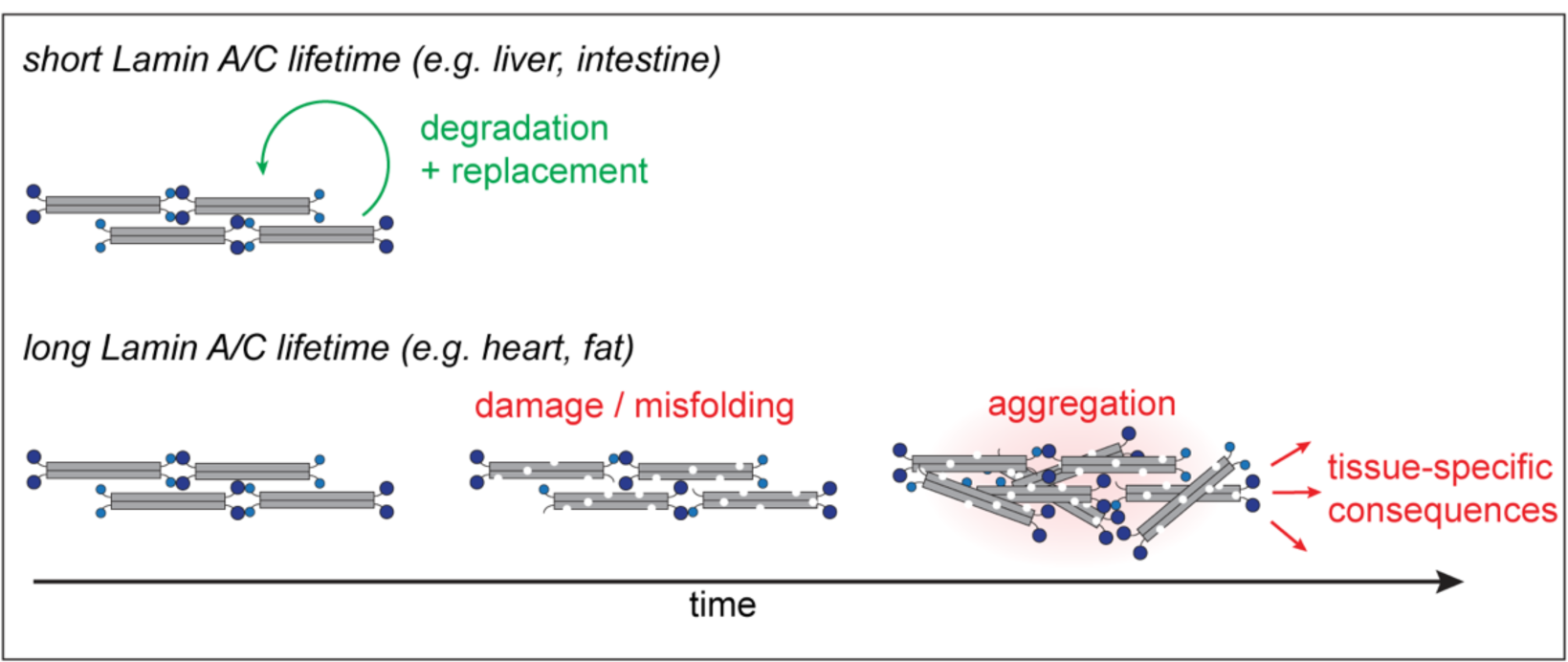
Working model of interplay between protein turnover rate and time- and mutation-dependent accumulation of dysfunction in progeroid tissues.

All gene therapies rely on a key assumption: that targeting a gene or transcript will lead to the desired changes to the encoded protein. We have demonstrated that Progerin is a long-lived and toxic mutant protein. Because long-lived proteins are rarely synthesized and rarely degraded, they are relatively insensitive to changes in their synthesis rate(*23*). This explains why gene therapies have significant latency and limited potency in decreasing Progerin protein levels in the cardiovascular system(*6*). Interestingly, combination treatment with a farnesyltransferase inhibitor, which interferes with the biosynthesis of Progerin at the protein level, results in more robust clearance of the mutant protein(*6*). Further exploration of orthogonal approaches for interfering with Progerin protein production or accelerating protein degradation may identify new therapies for this devastating disease.

By evaluating the relationship between human disease alleles and protein lifetime, we discovered that disease-linked proteins are generally over-represented in the long-lived proteome. As we have demonstrated in the context of HGPS, long protein lifetime may exert an important influence on protein function and present a major barrier to treatment in numerous human disease states.

## Materials and Methods

### Mouse model

All animal experiments were performed in compliance with relevant ethical regulations and with approval by the Institutional Animal Care and Use Committee at UCSF (IACUC protocol number AN178187, PI: A.B.).

### Protein extraction from mouse tissues for Western blotting

Approximately 30 milligrams of frozen tissue was excised on dry ice with a clean razorblade and placed in a fresh tube. 100 uL of urea lysis buffer (ULB: 8M urea, 75 mM NaCl, 50 mM HEPES pH 7.9, protease and phosphatase inhibitors) was added to the tube, and the tissue was then minced on ice with scissors. ULB was added to a final volume of 600 uL, and the sample was transferred to a Dounce homogenizer. The sample was homogenized for ∼40 strokes with the tight pestle, then transferred to a clean microcentrifuge tube. The sample was then probe sonicated at 4C (10% amplitude, 10 seconds, 2 cycles) before being centrifuged (21,000 x g, 11 minutes, 4C). The supernatant was transferred to a clean tube. Protein concentration was quantified by microBSA assay (Pierce). Lysates were mixed with SDS-PAGE sample buffer, heated to 95°C for 5 minutes, and cooled. 20 micrograms of protein were loaded per well for Western blot analysis and proteins were detected with an HRP-conjugated mouse monoclonal antibody that recognizes an N-terminal epitope shared in Lamin A, Lamin C, and Progerin (Santa Cruz sc-376248). Blots were visualized on a Chemi-Doc (BioRad) and proportional abundance of each protein isoform was determined by densitometry.

### Serial extractions of protein from mouse tissues

Relative solubility of the Lamin proteins was assayed using a serial extraction protocol adapted from (*30*). Approximately 20 milligrams of frozen tissue was excised on dry ice with a clean razorblade and placed in a fresh tube. Buffer 1 (10 mM HEPES pH 7.4, 2 mM MgCl2, 25 mM KCl, 250 mM sucrose, 1 mM DTT, protease and phosphatase inhibitors; 300 uL) was added to the tube, and tissue was minced on ice. The sample was transferred to a Dounce homogenizer and homogenized for ∼40 strokes with the tight pestle, then transferred to a clean microcentrifuge tube. The sample was probe sonicated at 4C (10% amplitude, 5 seconds, 2 cycles); a 50 uL aliquot (whole tissue lysate) was retained. The remaining sample was then centrifuged (20,000 x g, 5 minutes, 4C). The supernatant (S1) was retained. The pellet was resuspended in Buffer 2 (20 mM HEPES pH 7.4, 1 M NaCl, protease and phosphatase inhibitors; 250 uL) and incubated 20 minutes at RT with end-over-end rotation. The sample was then centrifuged (20,000 x g, 5 minutes, RT) and the supernatant (S2) was retained. The pellet was resuspended in Buffer 3 (20 mM HEPES pH 7.4, 1 M NaCl, 2% Triton-X-100, protease and phosphatase inhibitors; 250 uL) and incubation and centrifugation were repeated. The supernatant (S3) was retained and the pellet was resuspended in Buffer 4 (20 mM HEPES pH 7.4, 1M NaCl, 4M urea, protease and phosphatase inhibitors; 250 uL) and incubation and centrifugation were repeated. Finally, the supernatant (S4) was retained and the pellet was resuspended in Buffer 5 (20 mM HEPES pH 7.4, 1 M NaCl, 8M urea, protease and phosphatase inhibitors; 250 uL) and incubation and centrifugation were repeated. The final supernatant (S5) was retained. An aliquot of the tissue lysate and equal proportions of each supernatant were loaded on SDS-PAGE gels and processed for Western blotting with an HRP-conjugated mouse monoclonal antibody that recognizes an N-terminal epitope shared in Lamin A, Lamin C, and Progerin (Santa Cruz sc-376248).

### RNA extraction from mouse tissues

Approximately 30 milligrams of frozen tissue was excised on dry ice with a clean razorblade and placed in a fresh microcentrifuge tube. 100 uL of TRIzol was added, and tissue was minced with scissors on ice. Additional TRIzol was added to a final volume of 500 uL and the sample was transferred to a Dounce homogenizer. The sample was homogenized for ∼40 strokes with the tight pestle, then transferred to a clean microcentrifuge tube. The sample was incubated for 5 minutes at RT before chloroform (100 uL) was added. The sample was mixed by inversion and incubated for 3 minutes, then centrifuged (12,000 x g, 15 minutes, 4C). The aqueous RNA-containing supernatant was carefully removed and pipetted into a fresh tube. 1 uL of GlycoBlue and 250 uL isopropanol were added to the supernatant, incubated for 10 minutes at RT, then centrifuged (12,000 x g, 10 minutes, 4C). The supernatant was removed from the blue RNA-glycogen pellet, then the pellet was resuspended in 500 uL 75% ethanol and centrifuged (7500 x g, 5 minutes, 4C). The supernatant was removed and the pellet was allowed to air dry for 5-10 minutes. The pellet was resuspended in 50 uL nuclease-free water (Ambion) and RNA concentration was determined by NanoDrop.

### Transcript quantification with digital droplet PCR

For each sample, 1 ug of RNA was diluted into 15 uL reverse transcription reactions using the iScript RT kit (Bio-Rad) and reactions were carried out following the manufacturer’s instructions. Each cDNA reaction was diluted to 10 ng/uL (RNA equivalents). For each sample, 20 ng (RNA equivalents) was combined with 10 uL of 2x ddPCR Supermix for Probes (no UTP; Bio-Rad), 1 uL of a 20x FAM primer/probe set, 1 uL of a HEX primer/probe set, and nuclease free water to a final volume of 20 uL. *Lmna* primers and probe sequences are listed below and are adapted from(*6*). *mTfrc* primers and probe sequence was from Bio-Rad (PrimerPCR ddPCR Expression Probe Assay for Tfrc Mouse (HEX)). Reaction droplets were generated on a Droplet Maker (Bio-Rad); ddPCR reactions were run on a C1000 Thermocycler (Bio-Rad); and reactions were analyzed on a QX100 ddPCR reader (Bio-Rad) according to the manufacturer’s instructions. ddPCR allows the absolute quantification of transcript abundance based on the ratio of positive to negative droplets and is unaffected by primer-specific variables such as reaction efficiency(*44*). We used absolute ddPCR quantification to determine the abundance of the *Lmna*, *Lmnc,* and *Progerin* transcripts (copies per droplet). *mTfrc* reactions were run in the HEX channel of each ddPCR reaction as a positive control, but were not used to normalize *Lmna* transcript measurements. Transcript abundance was expressed as a proportion for each transcript. For example, for *Lmna*:

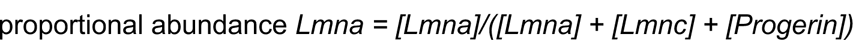

ddPCR probe and primersets for mouse *Lmna, Lmnc,* and *Progerin* were as previously described(*6*) and as listed in the following table.

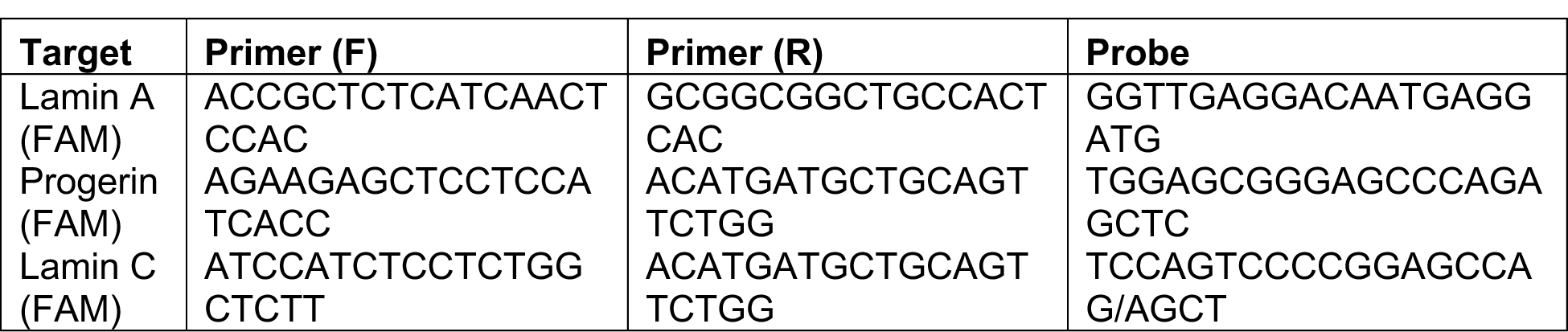

### Metabolic labeling of mice and tissue isolation

As recently described(*14*), we performed a 6-timepoint, 32-day ^15^N labeling timecourse (0, 2, 4, 8, 16, and 32 days of labeling) with a total of 3 animals of both sexes per labeled timepoint, and 2 animals for the day 0 (unlabeled) timepoint. Timecourses were performed in *LMNA^G609G/+^* C57Bl/6 mice at approximately 9 weeks of age. ^14^N and ^15^N mouse chow was obtained from Silantes. Animals were first habituated to the chow formulation by feeding ^14^N (normisotopic) food for 1 week; animals were then transitioned to ^15^N chow throughout the labeling period (roughly 3 grams / animal / day). Animals were sacrificed by CO_2_ inhalation followed by cervical dislocation, followed by tissue dissection and flash freezing by submersion in liquid nitrogen.

### Protein extraction and sample preparation for LC-MS/MS

#### Protein extraction

Approximately 30 milligrams of frozen tissue was excised on dry ice with a clean razorblade and placed in a fresh tube. 100 uL of protein extraction buffer (PEB: 5% SDS, 100 mM TEAB, protease and phosphatase inhibitors, pH ∼7) was added to the tube. The tissue was minced on ice; PEB was added to bring the final volume to 600 uL, then the sample was transferred to a Dounce homogenizer. The sample was homogenized for ∼40 strokes with the tight pestle, then transferred to a clean microcentrifuge tube. The sample was then probe sonicated at 4C (10% amplitude, 10 seconds, 2 cycles) before being centrifuged (21,000 x g, 11 minutes, 4C). The supernatant was transferred to a clean tube, and aliquots were separated for proteomics and protein quantification by microBSA assay (Pierce).

#### Trypsinization

Samples were diluted to 1 mg/mL in 5% SDS, 100 mM TEAB, and 25 µg of protein from each sample was reduced with dithiothreitol to 2 mM, followed by incubation at 55°C for 60 minutes. Iodoacetamide was added to 10 mM and incubated in the dark at room temperature for 30 minutes to alkylate proteins. Phosphoric acid was added to 1.2%, followed by six volumes of 90% methanol, 100 mM TEAB. The resulting solution was added to S-Trap micros (Protifi), and centrifuged at 4,000 x g for 1 minute. The S-Traps containing trapped protein were washed twice by centrifuging through 90% methanol, 100 mM TEAB. 1 µg of trypsin was brought up in 20 µL of 100 mM TEAB and added to the S-Trap, followed by an additional 20 µL of TEAB to ensure the sample did not dry out. The cap to the S-Trap was loosely screwed on but not tightened to ensure the solution was not pushed out of the S-Trap during digestion. Samples were placed in a humidity chamber at 37°C overnight. The next morning, the S-Trap was centrifuged at 4,000 x g for 1 minute to collect the digested peptides. Sequential additions of 0.1% TFA in acetonitrile and 0.1% TFA in 50% acetonitrile were added to the S-trap, centrifuged, and pooled. Samples were frozen and dried down in a Speed Vac (Labconco) prior to TMTpro labeling.

#### TMT labeling

Samples were reconstituted in TEAB to 1 mg/mL, then labeled with TMTpro 16plex reagents (Thermo Fisher) following the manufacturers protocol. Briefly, TMTpro tags were removed from the −20°C freezer and allowed to come to room temperature, after which acetonitrile was added. Individual TMT tags were added to respective samples and incubated at room temperature for 1 hour. 5% hydroxylamine was added to quench the reaction, after which the samples for each experiment were combined into a single tube. Since we performed abundance quantitation on unlabeled peptides, 0 day samples were added to four of the unused channels, increasing the signal for the unlabeled peptides. TMTpro tagged samples were frozen, dried down in the Speed Vac, and then desalted using homemade C18 spin columns to remove excess tag prior to high pH fractionation.

#### High pH Fractionation

Homemade C18 spin columns were activated with two 50 µL washes of acetonitrile via centrifugation, followed by equilibration with two 50 µL washes of 0.1% TFA. Desalted, TMTpro tagged peptides were brought up in 50 µL of 0.1% TFA and added to the spin column. After centrifugation, the column was washed once with water, then once with 10 mM ammonium hydroxide. Fractions were eluted off the column with centrifugation by stepwise addition of 10 mM ammonium hydroxide with the following concentrations of acetonitrile: 2%, 3.5%, 5%, 6.5%, 8%, 9.5%, 11%, 12.5%, 14%, 15.5%, 17%, 18.5%, 20%, 21.5%, 27%, and 50%. The sixteen fractions were concatenated down to 8 by combining fractions 1 and 9, 2 and 10, 3 and 11, etc. Fractionated samples were frozen, dried down in the Speed Vac, and brought up in 0.1% TFA prior to mass spectrometry analysis.

#### LC-MS/MS Analysis

*Data collection:* Peptides from each fraction were injected onto a homemade 30 cm C18 column with 1.8 um beads (Sepax), with an Easy nLC-1200 HPLC (Thermo Fisher), connected to a Fusion Lumos Tribrid mass spectrometer (Thermo Fisher). Solvent A was 0.1% formic acid in water, while solvent B was 0.1% formic acid in 80% acetonitrile. Ions were introduced to the mass spectrometer using a Nanospray Flex source operating at 2 kV. The gradient began at 3% B and held for 2 minutes, increased to 10% B over 7 minutes, increased to 38% B over 94 minutes, then ramped up to 90% B in 5 minutes and was held for 3 minutes, before returning to starting conditions in 2 minutes and re-equilibrating for 7 minutes, for a total run time of 120 minutes. The Fusion Lumos was operated in data-dependent mode, employing the MultiNotch Synchronized Precursor Selection MS3 method to increase quantitative accuracy(*45*). The cycle time was set to 3 seconds. Monoisotopic Precursor Selection (MIPS) was set to Peptide. The full scan was done over a range of 400-1500 m/z, with a resolution of 120,000 at m/z of 200, an AGC target of 4e5, and a maximum injection time of 50 ms. Peptides with a charge state between 2-5 were picked for fragmentation. Precursor ions were fragmented by collision-induced dissociation (CID) using a collision energy of 35% and an isolation width of 1.0 m/z. MS2 scans were collected in the ion trap with an AGC target of 1e4 and a maximum injection time of 35 ms. MS3 scans were performed by fragmenting the 10 most intense fragment ions between 400-2000 m/z, excluding ions that were 40 m/z less and 10 m/z greater than the precursor peptide, using higher energy collisional dissociation (HCD). MS3 ions were detected in the Orbitrap with a resolution of 50,000 at m/z 200 over a scan range of 100-300 m/z. The isolation width was set to 2 Da, the collision energy was 60%, the AGC was set to 1e5, and the maximum injection time was set to 100 ms. Dynamic exclusion was set to 45 seconds.

#### Data analysis

Raw data was searched using the SEQUEST search engine within the Proteome Discoverer software platform, version 2.4 (Thermo Fisher), using the Uniprot mouse database (downloaded January 2020). Trypsin was selected as the enzyme allowing up to 2 missed cleavages, with an MS1 mass tolerance of 10 ppm, and an MS2 mass tolerance of 0.6 Da. Carbamidomethyl on cysteine, and TMTpro on lysine and peptide N-terminus were set as a fixed modifications, while oxidation of methionine was set as a variable modification. Percolator was used as the FDR calculator, filtering out peptides which had a q-value greater than 0.01. Reporter ions were quantified using the Reporter Ions Quantifier node, with an integration tolerance of 20 ppm, and the integration method being set to “most confident centroid”. Protein abundances were calculated by summing the signal to noise of the reporter ions from each identified peptide, while excluding any peptides with an isolation interference of greater than 30%, or SPS matches less than 65%.

#### Kinetic model

The kinetic model applied in this study has been previously described(*46*). Briefly, we are assuming that protein synthesis is a zero order process, occurs at a constant fractional rate, and that that the total protein concentration of each cell does not change during the experimental time-course. The dilution of the protein pool due to cell division can be modeled as a first order exponential process. Thus, the fractional turnover of unlabeled proteins during the labeling time course can be regarded as a first order kinetic process that can be modelled based on the following exponential equation:

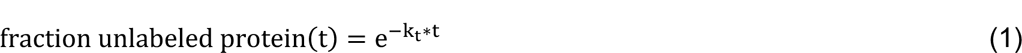

Where:

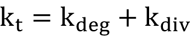

k_t_ is the clearance rate (observed rate of fractional labeling), k_deg_ is the rate of protein degradation and k_div_ is the rate of cell division.

The determination of k_t_ values were conducted as previously described(*46*) using the decay of the TMT reporter signals of unlabeled proteins. Protein-level TMT reporter abundances for unlabeled proteins for each replicate experiment were first normalized by dividing by the intensity of the t0 reporter and then the replicate experiments were aggregated in a single kinetic curve. In fitting the exponential decay curves of the unlabeled protein signals, a constant fractional baseline at infinite time was incorporated in the fitting equation. The equation used for fitting the curves was therefore: *intensity = baseline* + (1 - *baseline*)\**e*^−*k_t_*\**time*^. The goodness of fit for least squares fits were assessed by determining the R^2^, P-value and t-statistic of the fits (see Supp Table 1). For subsequent analyses, only protein k_t_ measurements that were obtained from all three replicate experiments, incorporated data from 4 or more peptide spectral matches (PSMs), and had t-statistic values greater than three were considered.

### Nucleic acid extraction and sample preparation for LC-MS/MS

#### Genomic DNA extraction

Approximately 30 milligrams of frozen tissue was excised on dry ice with a clean razorblade and placed in a fresh tube. 100 uL of TRIzol reagent (Invitrogen) was added and the tissue was rapidly minced on ice. An additional 400 uL of TRIzol was added, and the sample was then transferred to a Dounce homogenizer. The tissue was subjected to ∼40 strokes with the tight pestle until smooth, then transferred back to the original tube. The sample was incubated for at least 5 minutes before the addition of 100 uL chloroform followed by mixing and a further 3 minutes of incubation. The sample was then centrifuged (12,000 x g, 15 minutes, 4C) and the upper RNA-containing aqueous layer was discarded. 150 uL of absolute ethanol was added to the remaining sample, then inverted several times to mix. After 3 minutes of incubation at room temperature, the sample was centrifuged (2,000 x g, 5 minutes, 4C). The protein-containing supernatant was removed, then the DNA-containing pellet was resuspended in 500 uL of absolute ethanol and incubated for 30 minutes. The sample was then centrifuged (2,000 x g, 5 minutes, 4C), and the supernatant discarded. Sequential washes were then repeated with 95%, 85%, and 75% ethanol, after which the pellet was air-dried for 5-10 minutes. The pellet was then resuspended in 200 uL nuclease-free water (Ambion) at 56C, then incubated at 56C with shaking for 30 minutes to resuspend the pure DNA. The sample was centrifuged (12,000 x g, 10 minutes, 4C), then the supernatant containing pure DNA was moved to a clean tube. DNA concentration was determined with a NanoDrop spectrophotometer.

#### Digestion of genomic DNA to short oligonucleotides

3-5 micrograms of pure genomic DNA was diluted to a 50 uL volume in nuclease-free water, then combined with 50 uL of 2x Dinucleotide Buffer (DB: 5 mU/uL benzonase, 40 mU/uL shrimp alkaline phosphatase, 20 mM Tris pH 7.9, 100 mM NaCl, 20 mM MgCl2). Samples were incubated overnight at 37C. Spin-X UF Concentrators (Corning) were rinsed with 200 uL buffer (20 mM Tris pH 7.9, 100 mM NaCl, 20 mM MgCl2), then samples were applied and centrifuged through (12,000 x g, 5 min, RT). The eluate was collected for analysis.

#### Digestion of genomic DNA to mononucleosides

We extracted mononucleosides from genomic DNA similarly to a previously described method(*47*) with some modifications. 1-3 micrograms of pure genomic DNA was diluted to a 50 uL volume in nuclease-free water, then combined with 50 uL of 2x Mononucleoside Buffer (MB: 5 mU/uL benzonase, 40 mU/uL shrimp alkaline phosphatase, 60 uU/uL phosphodiesterase I, 20 mM Tris pH 7.9, 100 mM NaCl, and 20 mM MgCl2). Samples were incubated overnight at 37C. Spin-X UF Concentrators (Corning) were rinsed with 200 uL buffer (20 mM Tris pH 7.9, 100 mM NaCl, 20 mM MgCl2), then samples were applied and centrifuged through (12,000 x g, 5 min, RT). The eluate was collected for analysis.

### Mononucleoside and Dinucleotide LC-MS/MS

Mononucleoside analyses were carried out by adapting a previously described method(*48*) using a Dionex Ultimate 3000 UHPLC coupled to a Q Exactive Plus mass spectrometer (Thermo Scientific). After purification, analytes were separated on a Hypersil Gold 2.1 x 150 mm column, protected by a 2.1 x 10 mm Hypersil Gold guard column (Thermo Scientific). The mobile phases were A: 0.1% formic acid in water, and B: 0.1% formic acid in acetonitrile. The flow rate was set to 400 µL/min, and the column oven was set to 36°C. 10 µL of each sample was injected, and the analytes were eluted using the following gradient: 0 min- 0% B, 6 min- 0% B, 8.5 min- 80% B, 9.5 min- 80% B, 10 min- 0% B, 13 min- 0% B. The Q Exactive Plus was operated in positive mode with a heated electrospray ionization (HESI) source. The spray voltage was set to 3.5 kV, the sheath gas flow rate was set to 40, and the auxiliary gas flow rate set to 7, while the capillary temperature was set to 320°C. A parallel reaction monitoring (PRM) method was used to quantify the unlabeled nucleotide, along with all of its N15 isotopes in a single scan. This was accomplished by using wide (8 m/z) isolation widths when selecting the nucleotides for fragmentation. By employing this method, we were able to quantify the level of labeling by looking at the intensity of each N15 labeled base in the MS2 scan. Fragment ions were detected in the Orbitrap with a resolution of 70,000 at m/z 200. Using a high resolution MS2 scan allowed us to resolve N15 and C13 isotopes. Peak areas from the fragment ions were extracted with a 10 ppm mass tolerance using the LC Quan node of the XCalibur software (Thermo Scientific).

Dinucleotide analyses were carried out using the same instrumentation, column, mobile phases, column temperature, and flow rate employed by the mononucleoside experiments. The gradient was changed to optimize dinucleotide separation as follows: 0 min- 5% B, 0.5 min- 5% B, 2.5 min- 90% B, 3.25 min- 90% B, 3.5 min- 5% B, 5.5 min- 5% B. The Q Exactive Plus was operated using the same tune settings as the mononucleotide experiment. However, instead of a PRM method, a full scan method from 500-650 m/z was developed to quantify the dinucleotides dCdC, TT, dAdA, and dGdG, along with their corresponding N15 isotopes. Precursor ions were detected in the Orbitrap with a resolution of 140,000 at m/z 200, using the high resolution MS1 scan to try to separate N15 and C13 isotopes as much as possible. Peak areas from the fragment ions were extracted with a 10 ppm mass tolerance using the LC Quan node of the XCalibur software (Thermo Scientific).

#### Measurement of kdiv

To accurately measure rates of cell division (*k_div_*) while factoring in the effects of incomplete labeling and nucleotide recycling, we considered the time-dependent labeling patterns of mononucleosides derived from genomic DNA. Upon initiation of ^15^N labeling, newly synthesized DNA strands can incorporate nucleotides from a precursor pool that includes fully ^15^N-labeled nucleotides derived from the dietary source, partially labeled species (containing one to four ^15^N atoms) derived by biosynthesis from incompletely labeled ^15^N precursors, and completely unlabeled nucleotides derived from recycling. Therefore, it may not be possible to accurately determine the ratio of new to old strands (and hence *k_div_*) from the mononucleotide data alone. We previously used the labeling pattern of dinucleotides to resolve this ambiguity(*14*). The isotopologue distribution of labeled (non-monoisotopic) peaks in the dinucleotide spectra are dependent on the composition of the nucleotide precursor pool. Through regression analyses, we determined that within all tissues and timepoints analyzed in this study, the isotopologue distributions of the dinucleotide data could be best modeled based on the assumption that newly synthesized strands had very low levels of fully unlabeled nucleotides. Hence, the fractional population of labeled non-monoisotopic peaks within dinucleotide and mononucleotide data were consistent with each other and could be used to determine the fractional population of new strands. For each tissue, fractional labeling of mononucleotide and dinucleotides for all 4 bases were combined and the aggregated dataset was fit to a single exponential equation to determine first order rate constant for division (*k_div_*). These data appear in Table S2.

### Analysis of Proteomic Data

#### Quality filtering and analysis of k_t_ values

Proteomic data was acquired in the form of 16-plex TMT replicates containing 2 full 6-timepoint timecourses (one from wild type animals and one from progeroid animals). For each genotype within each TMT replicate, proteins were filtered to retain only those detected with at least 3 peptide spectral matches (PSMs) in all timepoints. Proteins that met these criteria were then filtered per genotype within each TMT replicate based on goodness of fit using the t-statistic. The t-statistic is equal to the turnover rate (*k_t_*) divided by the standard error of that value. This metric determines to what extent measurement error influences *k_t_*. We applied a minimum t-statistic cutoff of 3, meaning that the magnitude of the turnover rate *k_t_* is at least 3 times the magnitude of the standard error. Between 50% to 63% of detected proteins passed these coverage and goodness-of-fit criteria (Figure S2). Along with the sample size, the t-statistic can be used to determine a p-value that indicates the probability that the turnover rate reported has a meaningful non-zero value. The *k_t_,* standard error, t-statistic, and p-value for each protein are reported in Table S1. The *k_t_,* standard error, and sample size were used to perform per-protein statistical tests across tissues, to identify proteins with significantly different turnover kinetics between tissues. These data are reported in Table S1.

#### Determination of relative protein abundance within tissues

To evaluate relative protein abundance between genotypes within tissues of 3-month-old mice, technical replicate unlabeled wild type and progeroid samples from each multiplexed TMT run were first channel normalized, then the geometric mean was calculated to determine mean normalized intensities for each biological replicate. Protein abundance was then length-normalized by dividing each protein’s normalized intensity by the number of amino acids. Finally, samples were normalized for comparison across biological replicates by normalizing each channel to the maximum value detected. The geometric mean abundance across all biological replicates was calculated by determining the geometric mean of the length- and channel-normalized protein abundance. Finally, the fold change of protein abundances between wild type and progeroid tissues was determined.

Additional multiplexed TMT experiments were performed to compare protein abundances within tissues from 9-month-old wild type and progeroid mice. In these experiments, two biological replicate samples were analyzed in technical duplicate for each genotype in an 8-plex TMT experiment per tissue. Raw data were subjected to the same sequence of technical replicate averaging, protein length normalization, channel normalization, biological replicate averaging, and fold change determination described above.

#### Identification of disease-linked proteins

Disease associations were determined by cross-referencing proteomic datasets to the Online Mendelian Inheritance in Man (OMIM) database (www.omim.org). A complete list of OMIM allele – gene associations for the mouse genome was downloaded from OMIM, then cross-referenced to mouse proteins in each decile of turnover rate, ranging from 1^st^ decile (most long-lived) to 10^th^ decile (most short-lived) in each tissue. Over-representation of disease alleles in deciles was evaluated by Fisher’s exact test.

### Protein immunoprecipitation from mouse tissues

Metabolically labeled tissue extracts from progeroid animals prepared as described above in PEB for proteomic analysis were used for immunoprecipitation as follows. Approximately 1300-1500 ug of lysate (100 uL) was aliquoted into a fresh tube and diluted to 2 mL final volume in Dilution Buffer (10 mM Tris pH 7.4, 150 mM NaCl, 1% Triton-X-100, 1% deoxycholate, 2.5 mM MgCl2, protease and phosphatase inhibitors). For each sampe, 60 uL of Protein G magnetic beads (Life Technologies) were rinsed and resuspended in Dilution Buffer, then added to the diluted lysate. The sample was incubated at 4C with rotation for 30 minutes. Beads were removed and the pre-cleared supernatant was transferred to a clean tube; 5% of the sample was reserved as input. A mixture of two anti-Lamin A/C antibodies was used to immunoprecipitated Lamin A, Lamin C, and Progerin: Lamin A/C E-1 (Santa Cruz sc-376248, 20 uL per IP) and Lamin A/C (Active Motif #39288 Clone 3A6-4C11, 5 uL per IP). The IP was incubated overnight at 4C with nutation. 60 uL of pre-rinsed Protein G beads were then added to the sample and incubated for 30 minutes at 4C with nutation. Beads were collected on a magnetic stand and supernatant (flow-through) was saved in a fresh tube. Beads were washed in 3 x 1 mL of Wash Buffer (10 mM Tris pH 7.4, 150 mM NaCl, 0.5 mM EDTA, 0.1% SDS, 1% Triton-X-100, 1% deoxycholate, 2.5 mM MgCl2, protease and phosphatase inhibitors). Following the third wash, liquid was carefully removed using a gel loading tip; 2x SDS-PAGE sample buffer was added to the beads, and proteins were eluted by boiling for 5 minutes at 95C.

To check the efficiency of each IP, equal proportions of the input and flow-through fractions were analyzed by Western blotting for Lamin A/C (E-1 HRP, Santa Cruz, sc-376248 HRP). If a noticeable decrease in protein abundance was apparent in the flow-through sample, the IP samples were run on an SDS-PAGE gel and stained with Colloidal Coomassie stain. A gel slice containing the Lamin A, Lamin C, and Progerin bands was cut out for downstream mass spectrometry analysis.

### SIM/PRM-MS of Lamin and Progerin peptides

#### Data collection

SIM/PRM was performed as described above with the following changes. The gradient began at 3% B and held for 2 minutes, increased to 10% B over 5 minutes, increased to 38% B over 38 minutes, then ramped up to 90% B in 3 minutes and was held for 3 minutes, before returning to starting conditions in 2 minutes and re-equilibrating for 7 minutes, for a total run time of 60 minutes. The Fusion Lumos was operated using dual experiments, the first being a tSIM scan and the second being a PRM scan. Each tSIM scan targeted ions with an m/z of 597.7463 using an isolation window of 20 m/z, covering the range of the unique Progerin peptide and all possible N15 isotopes. The resolution was set to 120,000 at m/z of 200, an AGC target of 1e5, and a maximum injection time of 246 ms. The PRM scan fragmented precursor ions with an m/z of 589.7463 by collision-induced dissociation (CID) using a 1.6 m/z isolation width. The collision energy was set to 30% with a maximum injection time of 250 ms, and an AGC target of 1e4. The Ion Trap Scan Rate was set to ‘Rapid’.

#### Data analysis

The presence of Progerin was validated using raw MS2 data. Raw SIM data were searched for precursor ions with an m/z of 589.7463 and z of 2 in addition to all possible N15 isotopologues. The intensity of each isotopologue was measured using Thermo XCalibur Qual Browser (Table S5).

## Supporting information

Merged Supplementary Figures

## Acknowledgments

We would like to thank Yana Blokhina, Biao Wang, Esther Paolo Mirasol, Carlos Lizama Valenzuela, and Xi Chen for assistance with tissue dissections; Jonah Cool, Akiko Hata, and David Julius for critical reading of the manuscript; and all members of the Buchwalter lab for encouragement and ideas throughout the project period.

## Data and Materials availability

LC-MS/MS data have been deposited in the ProteomeXchange Consortium via the PRIDE partner repository, accessible at www.ebi.ac.uk/pride. The accession codes will be included in the final manuscript.

## Supplementary Materials

Materials and Methods

Figs. S1 to S6

Tables S1 to S5

