## Supplementary material for "Long lifetime and selective accumulation of the A-type lamins accounts for the tissue specificity of Hutchinson-Gilford progeria syndrome": Merged Supplementary Figures

Figure S1.

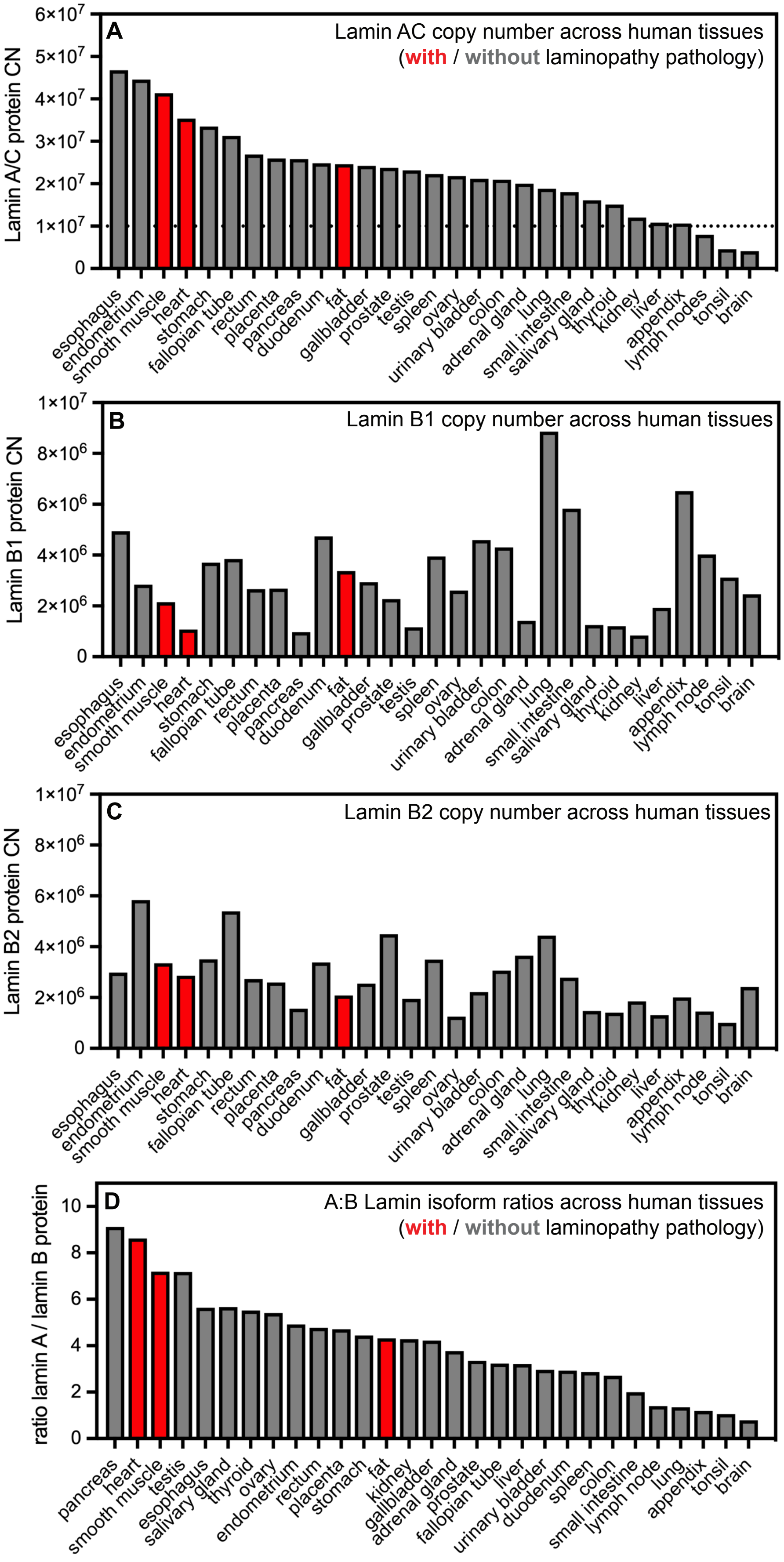


Figure S1. Estimated copy number per cell of Lamin A/C (A), Lamin B1 (B), and Lamin B2 (C) across human tissues, determined by histone normalization ("proteomic ruler" method). (D) Abundance ratio of A-type (Lamin A/C) to B-type (Lamin B1 and B2) lamin proteins across human tissues. All data reanalyzed from Wang et al *Mol Syst Biol* 2019.

Figure S2.

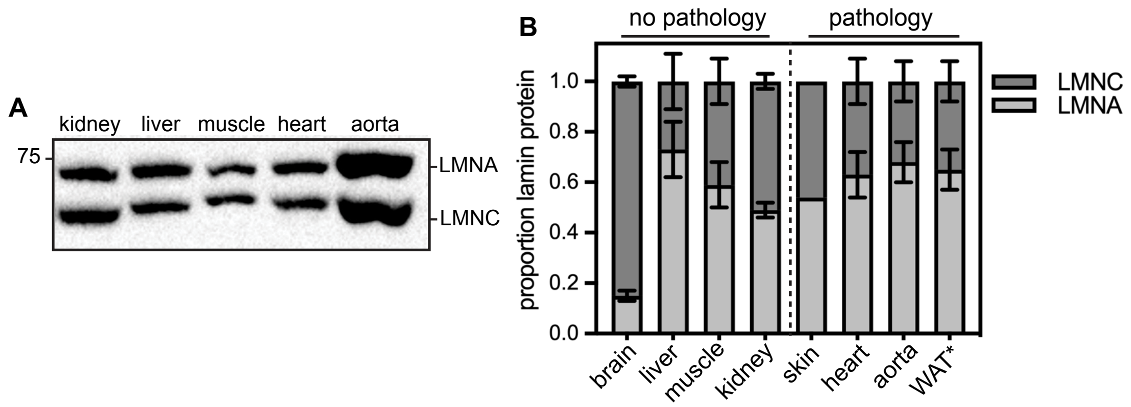


Figure S2. (A) Representative Western blot of tissue extracts from 9-month-old wild type mice. 20 ug of protein loaded per lane. (D) Quantification of proportional Lamin A and Lamin C protein isoform abundance by densitometry. * indicates that WAT was analyzed from 3 month old animals because of rapidly progressing lipodystrophy in these mice.

Figure S3. strong correlation between protein and RNA for SYK

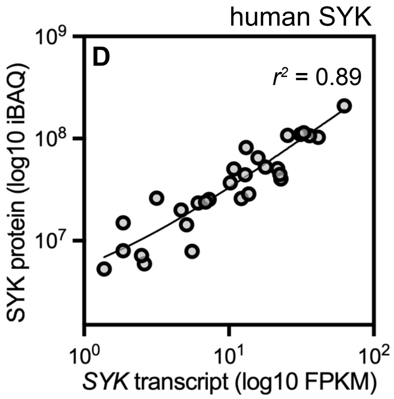


Figure S3. SYK protein abundance (iBAQ) and *SYK* RNA abundance (RNAseq FPKM) are strongly correlated ($r^2 = 0.89$) across 29 human tissues. Data reanalyzed from Wang et al., *Mol Syst Biol* 2019.

Figure S4

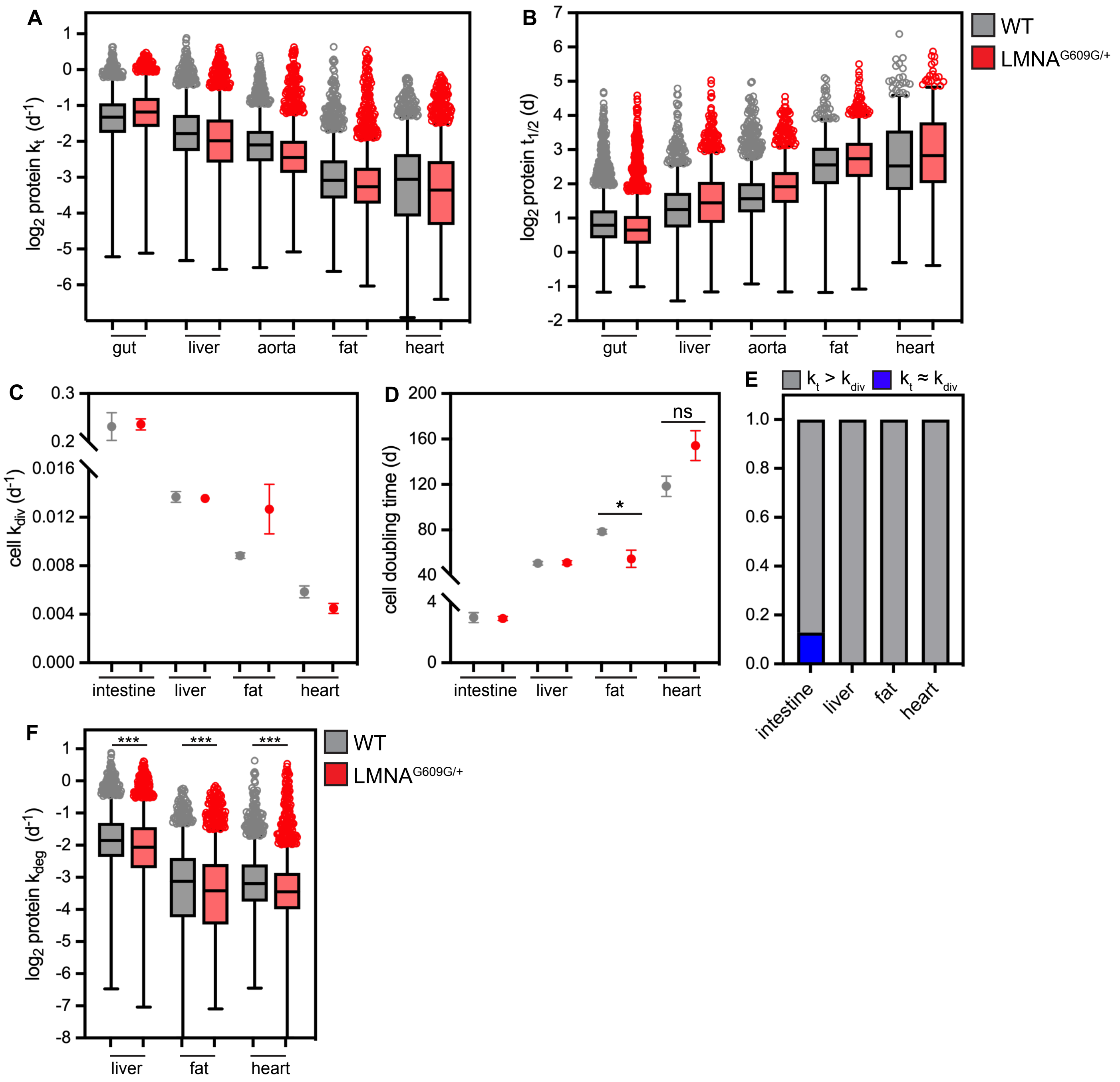


Figure S4. Protein turnover rates ($k_t$) (A) and predicted half-lives ($t_{1/2}$) (B) for proteins in tissues of healthy (gray) and progeroid (red) animals. Cell division rates ($k_{div}$) (C) and predicted cell doubling times (D) determined by TRAIL for tissues in healthy (gray) and progeroid (red) animals. * indicates that cells double significantly faster in progeroid fat. (E) Only the proliferative intestine has a significant number of proteins whose $k_t$ is equal to or less than $k_{div}$, suggesting that these proteins are diluted by cell division. (F) Protein degradation rates ($k_{deg}$) determined for proteins in liver, fat, and heart muscle of 3-month-old healthy and *LMNA*$^{G609G/+}$ mice. **** indicates a significant decrease in protein turnover flux in progeroid tissues.Data from wild type animals reproduced from Hasper et al., *Mol Syst Biol* 2023.

Figure S5.

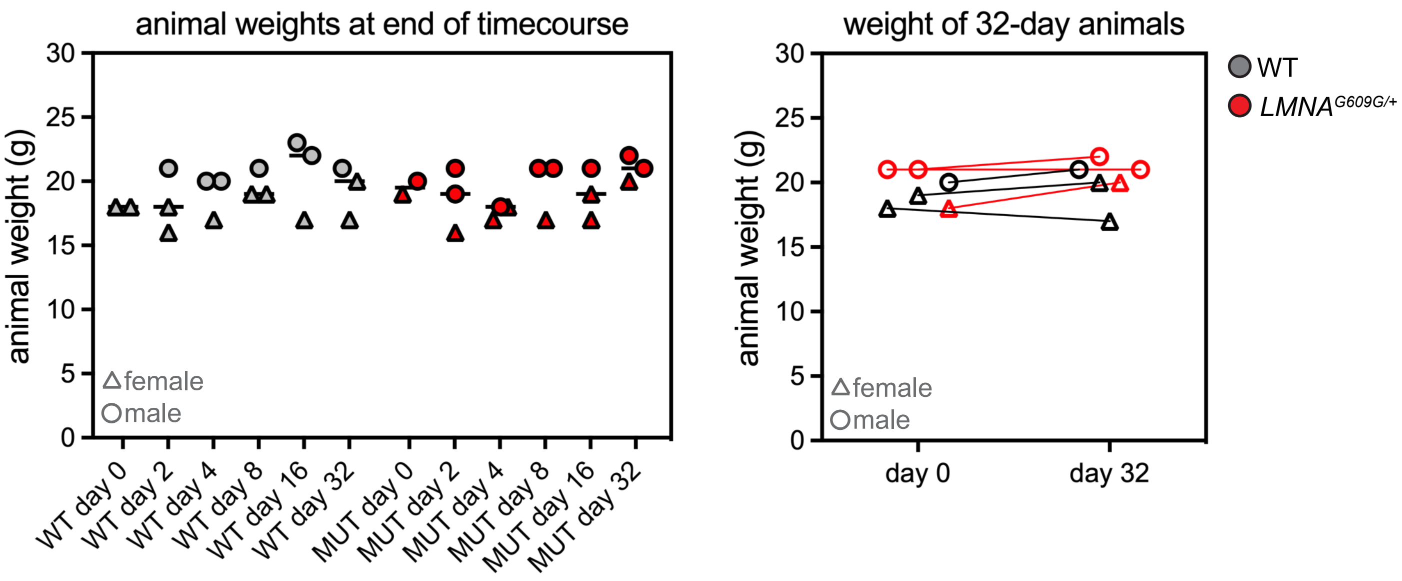


Figure S5. (A) Balanced mixtures of male and female mice were used for labeling timecourse. Weights of animals used for all timepoints of metabolic labeling are consistent. (B) Comparison of weights of same animals at beginning and end of 32-day timecourse shows no change.

Figure S6.

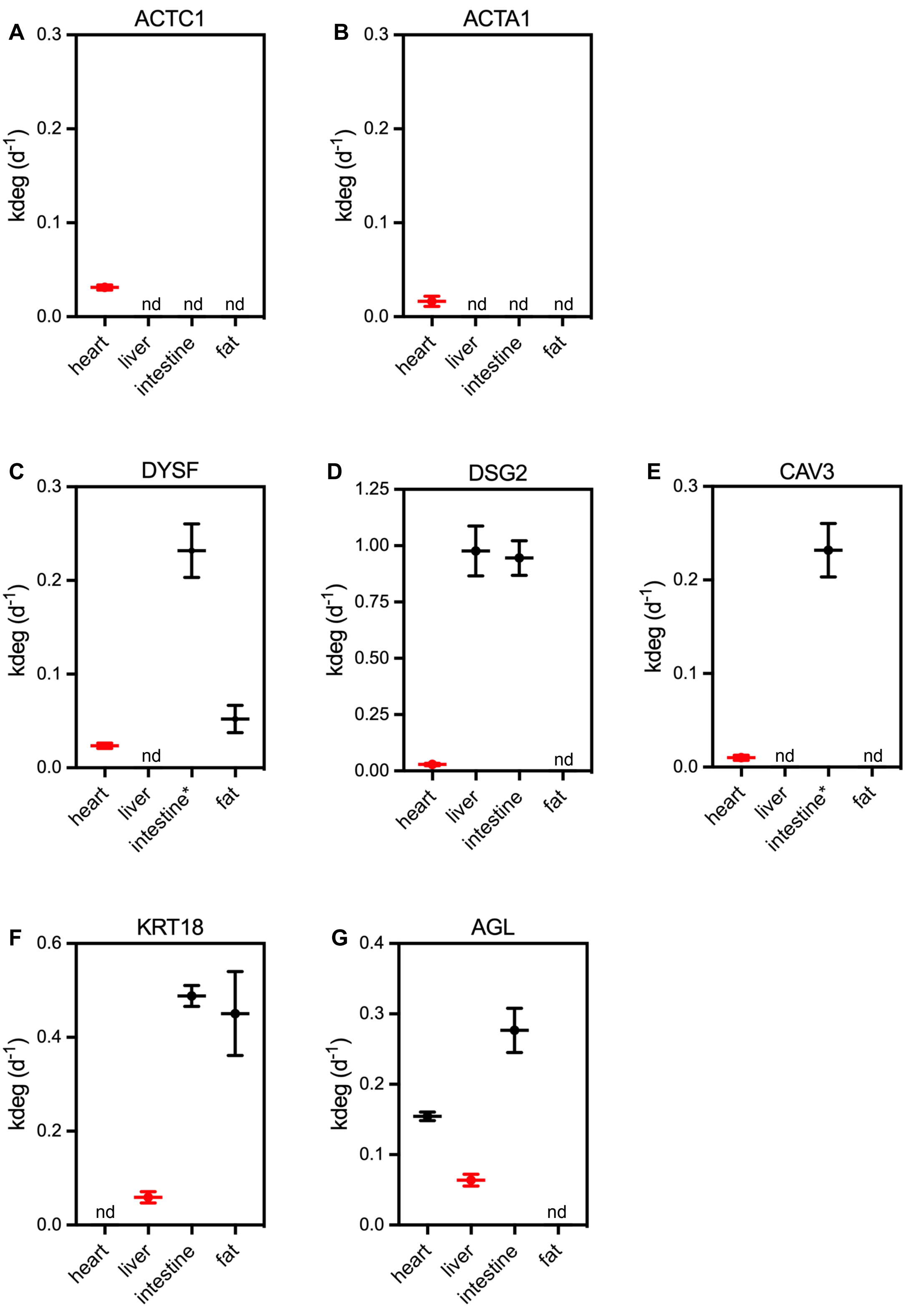


Figure S6. Turnover rates of selected proteins linked to human disease. (A-B) ACTC1 and ACTA1 are heart-specific long-lived proteins. (C-E) DYSF, DSG2, and CAV3 are broadly expressed but linked to cardiomyopathy, and are only long-lived proteins in the heart. (F-G) KRT18 and AGL are broadly expressed but linked to liver disease, and are only long-lived proteins in the liver.

**Table S3. turnover rate constants and predicted half-lives for LA/C/Progerin**

| tissue | genotype | $k_t$ ($d^{-1}$) | $k_{deg}$ ($d^{-1}$) | $t_{1/2}$ (days) | $t_{1/2corr}$ (days) |
|---|---|---|---|---|---|
| gut | wildtype | 0.146 ± 0.011 | ≈ kdiv (0.232) | 4.7 | 3.0# |
| | LMNA$^{(G609G/+)}$ | 0.110 ± 0.013 | ≈ kdiv (0.237) | 6.3 | 2.9# |
| liver | wildtype | 0.078 ± 0.012 | 0.064 ± 0.012 | 8.9 | 10.8 |
| | LMNA$^{(G609G/+)}$ | 0.077 ± 0.008 | 0.063 ± 0.021 | 9.0 | 11.0 |
| *aorta* | *wildtype* | 0.062 ± 0.006 | *nd* | 11.2 | *nd* |
| | *LMNA*$^{(G609G/+)}$ | 0.051 ± 0.004 | *nd* | 13.6 | *nd* |
| fat | wildtype | 0.041 ± 0.005 | 0.032 ± 0.005 | 16.9 | 21.7 |
| | LMNA$^{(G609G/+)}$ | 0.021 ± 0.008 | 0.008 ± 0.01 | 33.0 | 86.6* |
| heart | wildtype | 0.031 ± 0.003 | 0.026 ± 0.003 | 22.4 | 26.7 |
| | LMNA$^{(G609G/+)}$ | 0.027 ± 0.003 | 0.022 ± 0.003 | 25.7 | 31.5 |
